# Central thalamus modulates consciousness by controlling layer-specific cortical interactions

**DOI:** 10.1101/776591

**Authors:** Michelle J. Redinbaugh, Jessica M. Phillips, Niranjan A. Kambi, Sounak Mohanta, Samantha Andryk, Gaven L. Dooley, Mohsen Afrasiabi, Aeyal Raz, Yuri B. Saalmann

## Abstract

Consciousness is the capacity to experience one’s environment and internal states. The minimal mechanisms sufficient to produce this experience, the neural correlates of consciousness (NCC), are thought to involve thalamocortical and intracortical interactions, but the key operations and circuit paths are unclear. We simultaneously recorded neural activity in central thalamus and across layers of fronto-parietal cortex in awake, sleeping and anesthetized macaques. Spiking activity was selectively reduced in deep cortical layers and thalamus during unconsciousness, as were intracolumnar and interareal interactions at alpha and gamma frequencies. Gamma-frequency stimulation, when focused on the central lateral thalamus of anesthetized macaques, counteracted these neural changes and restored consciousness. These findings suggest that the NCC involve both corticocortical feedforward and feedback pathways coordinated with intracolumnar and thalamocortical loops.

**Summary:** Stimulation of central lateral thalamus counters anesthesia to restore wake cortical dynamics and consciousness.

## Main Text

Information processing during wakefulness involves feedforward pathways carrying sensory information from superficial layers to superficial/middle layers of higher-order cortical areas, and feedback pathways carrying priorities and predictions from deep layers to superficial or deep layers of lower-order cortical areas (*1, 2*). Information processing is altered during sleep, anesthesia and disorders of consciousness, though reported effects on feedforward (*3, 4*) and feedback (*5–7*) pathways have varied. Intracolumnar and thalamocortical interactions can influence feedforward and feedback pathways and promote recurrent processing (*8, 9*), and both intracolumnar (*10, 11*) and thalamocortical (*12, 13*) changes occur during unconsciousness. The central lateral nucleus (CL), of intralaminar thalamus, provides input to superficial layers and reciprocally connects with deep layers of fronto-parietal cortex (*14, 15*). Thus, CL is well-positioned to influence information flow between cortical layers and areas. CL damage is linked to disorders of consciousness (*14*), and deep brain stimulation of central thalamus increased responsiveness in a minimally conscious patient (*15*), though the mechanism is unknown. Resolving contributions of corticocortical and thalamocortical pathways to the NCC requires simultaneous recordings across layers in interconnected cortical areas and thalamus during different conscious states. Based on its connectivity, we hypothesized that CL influences feedforward, feedback and intracolumnar cortical processes to regulate information flow, and thereby consciousness.

Using linear multi-electrode arrays, we recorded spikes and local field potentials (LFPs) simultaneously from the right frontal eye field (FEF), lateral intraparietal area (LIP) – fronto-parietal areas implicated in awareness (*16*) – and interconnected CL in two macaques during task-free wakefulness, NREM sleep and general anesthesia (isoflurane, propofol). After characterizing thalamocortical network activity under anesthesia, we electrically stimulated the thalamus, inducing arousal. We evaluated signs of arousal before, during, and after stimulation using a customized scale similar to clinical metrics (supplementary materials), and performed statistical analyses using general linear models (Table S1-10).

Across 261 stimulation blocks, thalamic stimulation significantly increased arousal relative to pre- (F = 119.28, N = 261, p < 1.0×10^-10^) and post-conditions (F= 124.64, N = 261, p = 1.0×10^-10^) even accounting for differences in dose and anesthetic (Fig. 1, A and B; Fig. S1, A-C). Behavioral changes (Fig. 1A) were time-locked to stimulation: monkeys opened eyes with wake-like occasional blinks, performed full reaches/withdrawals with forelimbs (ipsi- or contralateral), made facial/body movements, showed increased reactivity (palpebral reflex, toe-pinch withdrawal) and altered vital signs (respiration rate, heartrate). Reconstruction of electrode tracks placed effective stimulations (arousal score ≥ 3) near CL center (Fig. 1, C-F). Euclidian proximity of the stimulation array to CL significantly predicted changes in arousal (Fig. S1, G-I; T = −3.39, N = 225, p = 0.00082); when systematically varying array depth, proximity to CL center showed a significant quadratic relationship with arousal (T = −2.92, N = 225, p = 0.00393; Fig. 1F; Fig. S1, D-F). Effective stimulation sites remained so on separate recording days and with different anesthetics (Fig. 1G). Importantly, stimulation effectiveness depended on frequency (Fig. 1, G and H). At effective sites, only 50 Hz stimulations reliably increased arousal (T = 3.91, N = 44, p = 0.00035). These results show that CL stimulation can rouse animals from stable, anesthetized states. This allowed us to zero-in on NCCs, here identified as activity differences between wakefulness and anesthesia which are selectively restored by effective (arousal score ≥ 3, N = 55, M = 4.70, SD = 1.70) relative to ineffective (arousal score < 3, N = 171, M = .77, SD = 0.74) 50 Hz stimulations.

**Fig. 1.**
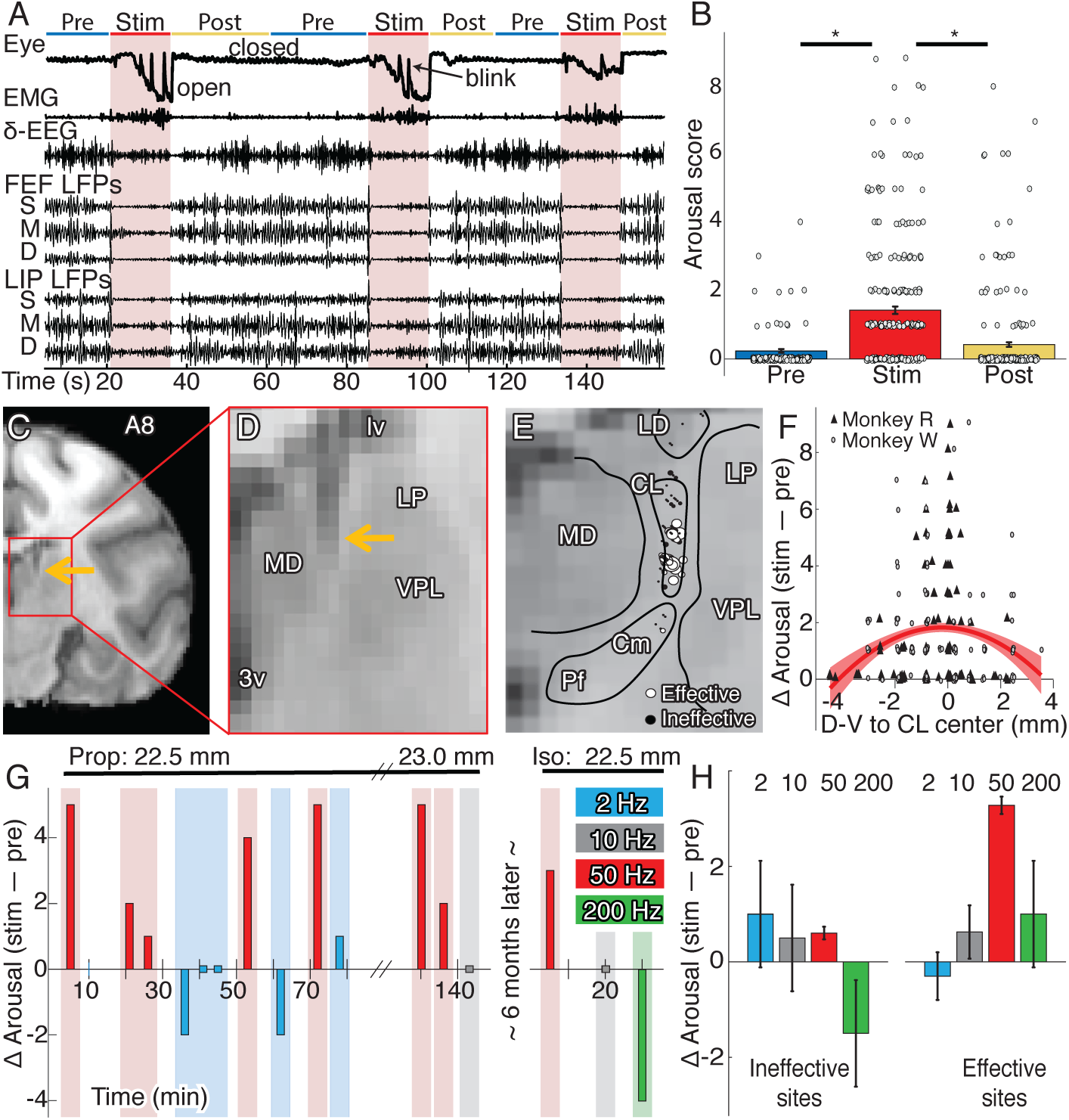
Gamma-frequency CL stimulation increased consciousness level. (**A**) Example behavioral and neural recordings during 50 Hz stimulation (arousal score 5). (**B**) Population mean arousal score (±SE) before, during and after stimulation (circles show individual stimulation events). (**C**) Coronal section of right hemisphere 8mm anterior to interaural line (A8). Arrow shows electrode. (**D**) Zoomed-in view of thalamus. (**E**) Stimulation sites in monkey R (N = 90) collapsed along A-P axis. Circles represent middle contact in stimulation array; diameter scales with induced arousal. (**F**) Stimulation-induced arousal change (score during stim – pre) as function of dorsal-ventral distance from CL center. Symbols show stimulation events by monkey; red curve shows quadratic fit (±SE). (**G**) Example stimulation series for different frequencies during propofol (left) and isoflurane (right) at same site 22.5 mm ventral to cortical surface. (H) Population mean arousal change (±SE of point estimate) for different stimulation frequencies at effective and ineffective sites.

We recorded 845 neurons across three brain areas (FEF, LIP, CL) during four states (wake, sleep, isoflurane, propofol; Fig. 2). Wake and anesthesia data derived from separate sessions, whereas the same neurons yielded sleep and wake data. Thalamic neurons showed state-dependent spike rate and bursting activity (Fig. 2, C and D). CL neurons recorded during anesthesia (T= −4.67, N = 282, p = 3.0*10^-5^) and NREM sleep (F = 16.40, N = 83, p = 0.001) had significantly lower spike rate than during wakefulness. Isoflurane and propofol effects were not significantly different (Fig. S2D). Relative to wakefulness, CL neurons also increased bursting during anesthesia (T = 2.27, N = 172, p = 0.024) and sleep (F = 7.11, N = 121, p = 0.0095).

**Fig. 2.**
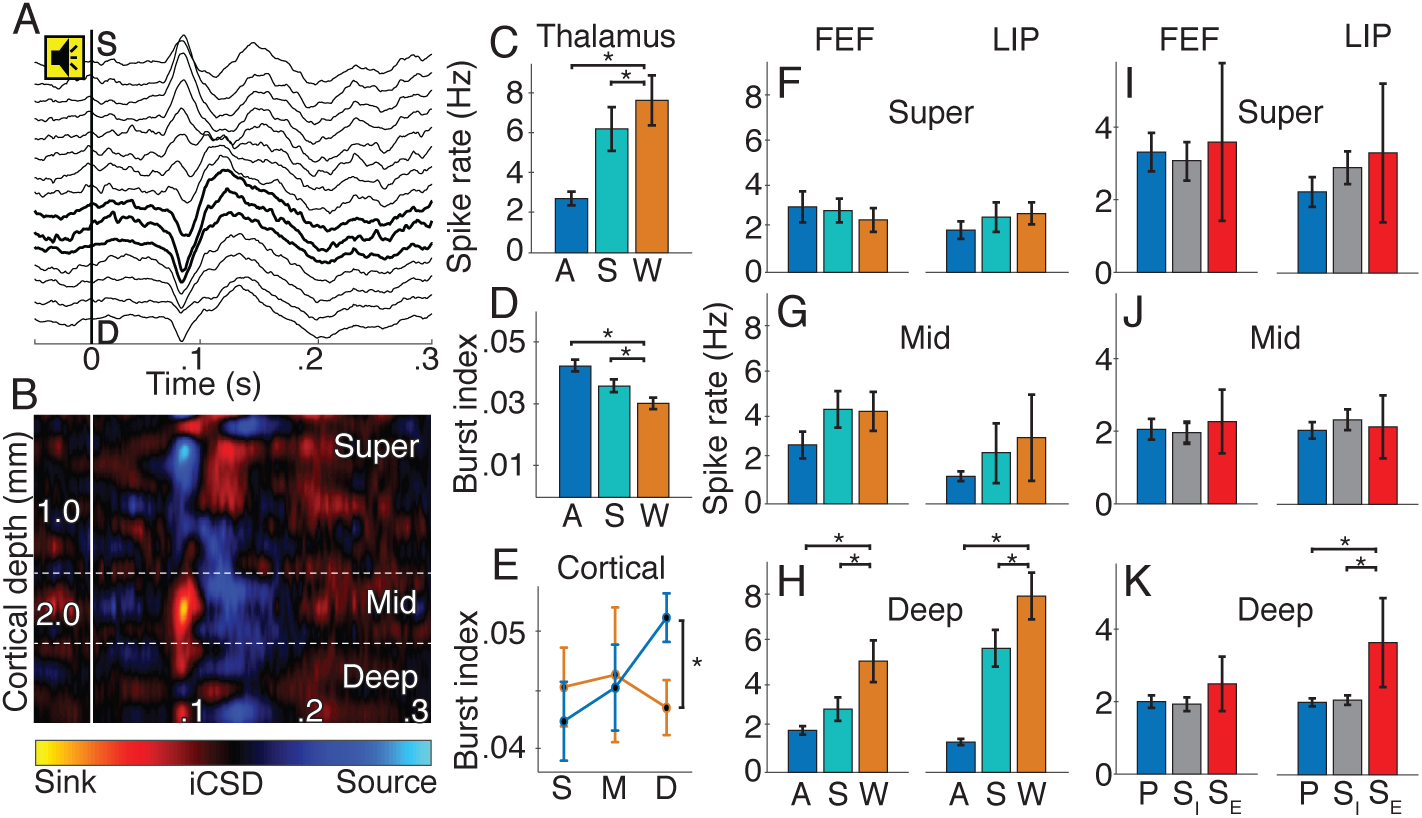
Consciousness level modulated spike rate and timing in deep cortical layers and CL. (**A**) Example sound-aligned evoked potentials from linear multielectrode array in FEF. Tone onset at 0 s. Bold lines show iCSD-defined middle layers. (**B**) Sound-aligned iCSD corresponding to A. (**C**) Population CL spike rate (±SE) and (**D**) CL burst index (±SE) for anesthesia (blue), sleep (teal), and wake (orange) states; * < 0.05. (**E**) Cortical (FEF and LIP) burst index (±SE) in superficial (S), middle (M) and deep (D) layers for wakefulness (orange) and anesthesia (blue). (**F**) Superficial, (**G**) middle and (**H**) deep layer spike rates (±SE) in FEF and LIP across states. (**I**) Superficial, (**J**) middle and (**K**) deep spike rates (±SE) in FEF and LIP during effective stimulation (S_E_, red), ineffective stimulation (S_I_, gray) and pre-stimulation (P, blue).

We localized cortical neurons to superficial, middle or deep layers using current source density (CSD) responses to sounds in the passive oddball paradigm (Fig. 2, A and B). Only deep neurons showed state-dependent activity (Fig. 2, E-H). Firing rates during sleep were significantly lower than wakefulness; the state by layer interaction was significant in both FEF (F = 15.17, N = 101, p = 0.008) and LIP (F = 7.70, N = 98, p = 0.031). Similarly, firing rates during anesthesia were lower than wake; state by layer interactions in FEF (T = 3.05, N = 281, p = 0.013) and LIP (T = 3.79, N = 282, p = 0.001) were significant. Only deep neurons increased bursting during anesthesia, evidenced by significant state by layer interaction (Fig. 2E; T = 2.12, N = 285, p = 0.035). Isoflurane and propofol yielded similar results (Fig. S2, A-C). Effective 50 Hz thalamic stimulation countered anesthesia effects in deep cortical layers of LIP (Fig 2, I-K); the four-way interaction of stimulation epoch, effectiveness, layer and area was significant (F = 5.19, N = 167, p = 0.023). Overall, states with higher consciousness level (stimulation-induced arousal, wake) showed increased deep cortical and thalamic activities, suggesting a key role in the NCC.

To measure state-related changes in thalamocortical and corticocortical communication, we calculated power and coherence using bipolar derivatized LFPs. We combined data across anesthetics, as effects were qualitatively similar (Fig. S3 and Fig. S4). We first focus on intracolumnar changes, particularly coherence within superficial layers, deep layers, and between superficial and deep layers of the same cortical area. Coherence changed markedly between wakefulness and anesthesia; anesthesia increased delta coherence (<4 Hz) and reduced alpha (8-15 Hz) and low gamma coherence (30-60 Hz; Fig. 3, A and B; Table S1 for complete statistics). Wake-anesthesia differences were consistent between different layers of FEF and LIP (Fig. 3 C-H), and qualitatively similar to differences between wakefulness and NREM sleep (Fig. S5). Notably, coherence between superficial and deep layers of both cortical areas showed substantial decreases in all higher-frequency (>4 Hz) communication during anesthesia (T ≥ 10.05, N = 8725, p < 1.0×10^-10^; Fig. 3, E and F; Table S1), suggesting altered processing in cortical microcircuits.

**Fig. 3.**
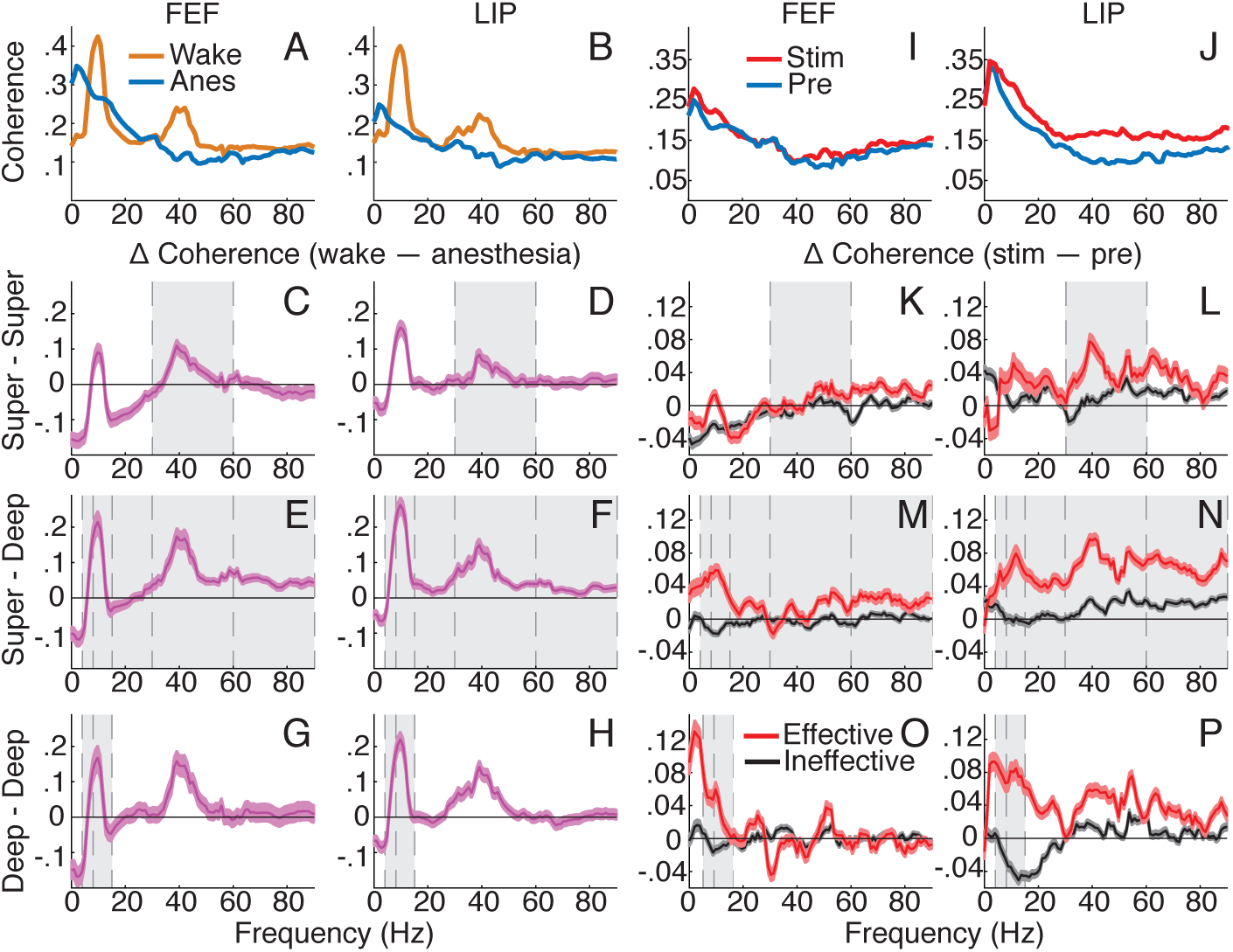
Intracolumnar interactions showed layer-specific NCC. (**A**) Population FEF and (**B**) LIP coherence ±SE (line thickness) for wakefulness and anesthesia. Average of all contact pairs across layers. (**C-H**) Population coherence difference between wakefulness and anesthesia. Positive when wake > anesthesia. Error bars indicate ±SE of T-tests at each frequency. Gray shading shows effects consistent between state (wake vs anesthesia) and thalamic stimulation (effective vs ineffective in K-P) results. Average of all contact pairs for/between: superficial (**C**) FEF and (**D**) LIP; superficial and deep (**E**) FEF and (**F**) LIP; deep (**G**) FEF and (**H**) LIP. (**I**) Population FEF and (**J**) LIP coherence ±SE under anesthesia before and during effective stimulation. Average across all layers. (**K-P**) Population coherence difference (stim – pre) ±SE for effective and ineffective stimulations. Positive when stim > pre. Average of all contact pairs for/between: superficial (**K**) FEF and (**L**) LIP; superficial and deep (**M**) FEF and (**N**) LIP; deep (**O**) FEF and (**P**) LIP.

Effective 50 Hz thalamic stimulation increased intracolumnar coherence differentially across frequency bands and layers (Fig. 3, K-P; Table S2 for complete statistics). Coherence within superficial layers increased for effective more than ineffective stimulations at low gamma (Fig. 3, K and L), showing a significant interaction between stimulation and effectiveness (T = 5.24, N = 2387, p =1.8×10^-6^). Within deep layers (Fig. 3, O and P), similar interactions show effective stimulations selectively increased coherence at theta (4-8 Hz, T = 9.04, N = 2183, p < 1.0×10^-10^) and alpha (T = 11.79, N = 2183, p < 1.0×10^-10^) frequencies. Importantly for intracolumnar processing, superficial and deep layers showed broadband coherence increases selective to effective stimulations at the same frequencies hindered by general anesthesia (bands > 4 Hz; T ≥ 4.83, N = 2631, p ≤ 1.3×10^-6^, see Table S2). Note that power changes during thalamic stimulation did not correlate with coherence changes (Fig. S6; Table S3 and S4 for complete statistics), thus power is neither a key contributor to the NCC nor stimulation-induced changes in coherence.

We next focus on anatomically-motivated interactions across fronto-parietal cortex: we measured coherence between the origin and termination of putative feedforward (superficial LIP-superficial/middle FEF) and two feedback (deep FEF- superficial LIP or deep FEF-deep LIP) pathways (*1, 2*). We also examined state-dependent effects on thalamocortical coherence (CL-superficial or CL-deep cortical layers) (*17, 18*). Communication between cortical areas showed significant changes during general anesthesia (Fig. 4A). Corticocortical coherence increased at delta and decreased at all higher frequencies across putative feedforward and feedback pathways (|T| ≥ 6.04, N ≥ 4030, p ≤ 1.6 x10^-9^; Table S5 for complete statistics). We found qualitatively similar effects during NREM sleep (Fig. S7) and for spike-field coherence (Fig. S8; Table S6 for complete statistics). Coherence between thalamus and either superficial or deep cortical layers decreased across all frequency bands during anesthesia (Fig. 4, I-L; Table S7 for complete statistics). Thalamocortical spike-field coherence showed similar effects (Fig. S9; Table S8 and Table S9 for complete statistics). These results show anesthesia decreases broadband thalamocortical and corticocortical coherence.

**Fig. 4.**
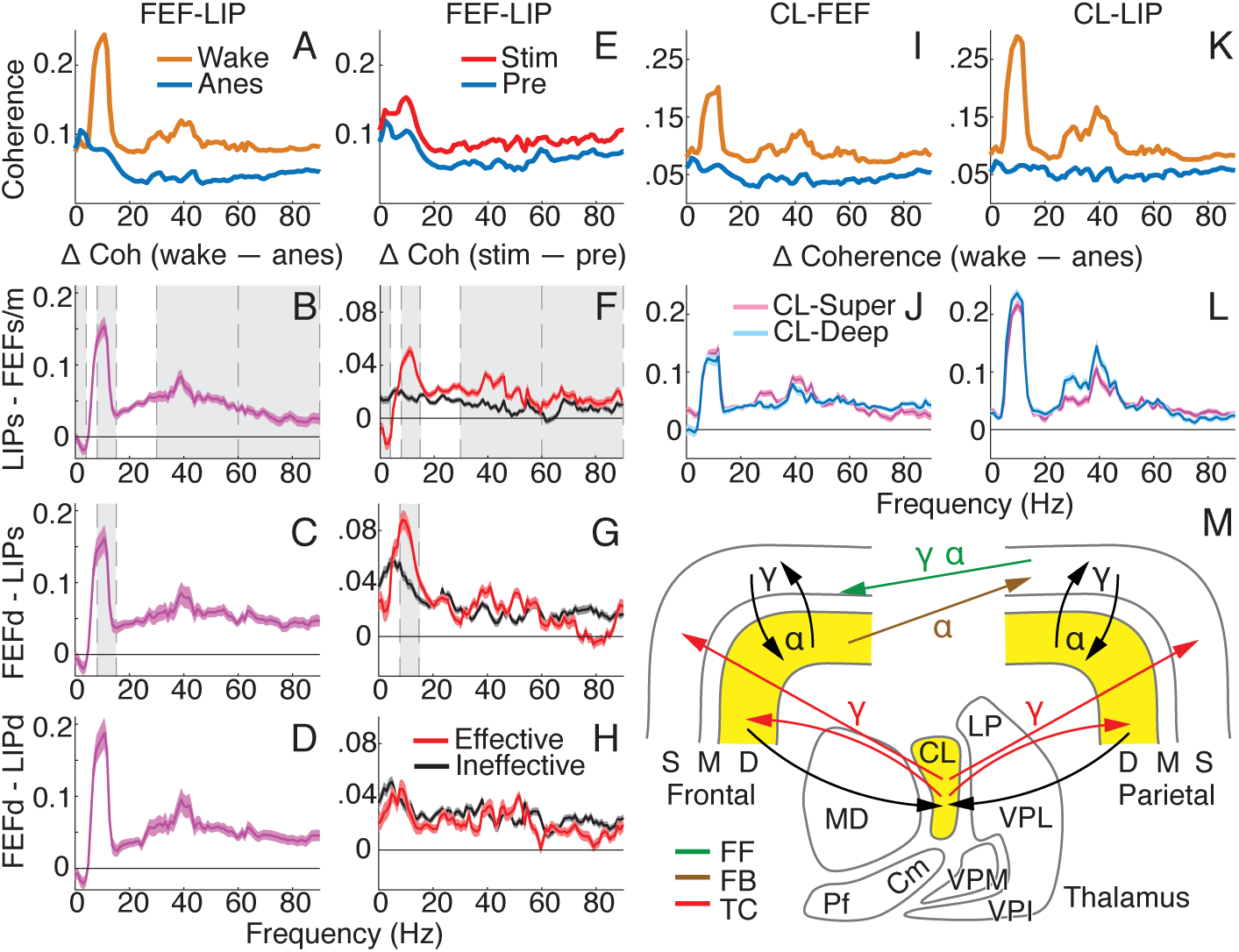
Thalamocortical and corticocortical interactions showed pathway-specific NCC. (**A**) Population coherence ±SE (line thickness) for all paired contacts between FEF and LIP during wakefulness and anesthesia. (**B-D**) Population average coherence difference: wake – anesthesia. Error bars indicate ±SE of T-tests at each frequency. Gray shading shows effects consistent between state and thalamic stimulation (in F-H) results. (**B**) Superficial LIP and superficial/mid FEF; (**C**) deep FEF and superficial LIP; (**D**) deep FEF and deep LIP. (**E**) Population coherence between FEF and LIP ±SE for all paired contacts under anesthesia, before and during effective stimulation. (**F-H**) Population average coherence difference, stim – pre, ±SE for effective and ineffective stimulations. (**F**) Superficial LIP and superficial/mid FEF; (**G**) deep FEF and superficial LIP; (**H**) deep FEF and deep LIP. (**I** and **K**) Population thalamocortical coherence ±SE for wake and anesthesia across all paired CL-FEF (**I**) and CL-LIP (**K**) contacts. (**J** and **L**) Population average thalamocortical coherence difference, wake – anesthesia, ±SE of T-tests. CL- superficial and CL-deep layers for (**J**) FEF and (**L**) LIP. (**M**) Schematic showing pathways and predominant frequencies contributing to NCC. Yellow shading where spiking changes with consciousness level. FF, feedforward; FB, feedback; TC, thalamocortical.

Thalamic stimulation isolated specific interactions between cortical areas important for consciousness (Fig. 4E). Effective stimulations resulted in targeted restoration of fronto-parietal coherence in putative feedforward and feedback pathways (Fig. 4, F-H). Coherence between superficial layers of LIP and superficial/middle FEF substantially reduced at delta (T= −4.05, N = 2799, p = 8.6×10^-4^), and increased at alpha (T = 6.87, N = 2799, p = 1.46×10^-10^), low gamma (T = 4.45, N = 2799, p = 1.50×10^-4^) and high gamma (60-90 Hz; T = 3.03, N = 2799, p = 0.027), for effective more than ineffective stimulations, as shown by significant interactions (Fig. 4F). There was also a significant interaction for coherence between deep layers of FEF and superficial LIP at alpha (Fig. 4G; T = 3.97, N = 1617, p = 0.001), showing substantial increases in alpha coherence specific to effective stimulations. While coherence between FEF and LIP deep cortical layers did generally increase with stimulation, no interactions were significant (Fig. 4H; Table S10 for complete statistics). Overall, more-conscious states showed increased alpha and gamma coherence in feedforward pathways as well as alpha coherence in the feedback pathways originating in deep layers and terminating in superficial layers of the lower-order area.

Our study suggests that both feedforward and feedback corticocortical pathways as well as intracolumnar and thalamocortical circuits generally contribute to the NCC (Fig. 4M), consistent with theories of consciousness that highlight feedforward, feedback and/or recurrent processing (*8, 9, 19, 20*). Specifically, we link consciousness to increased spiking activity in deep cortical layers and CL – consistent with cat/rodent studies of V1 (*21*) and CL (*22*) during NREM sleep – likely sustained through reciprocal deep cortex-CL connections. The deep cortical layers are anatomically positioned to drive feedback to superficial layers in lower-order areas, and to influence feedforward pathways via interactions with superficial layers in the cortical column. CL, with projections both to superficial and deep cortical layers, can modulate intracolumnar and cross-area interactions, which predominantly operated at alpha and gamma frequencies during consciousness. Consciousness thus seems particularly sensitive to disturbance of the deep cortex-CL loop, evident from our anesthesia/sleep effects. Reactivation of this loop with gamma-frequency CL stimulation reinstates cortical dynamics and increases consciousness level.

Several properties differentiate CL from other thalamic nuclei and make CL a suitable contributor to the NCC. CL receives input from the reticular activating system (*14*), and with projections to superficial and deep layers across fronto-parietal cortex, serves as an influential hub for network integration. Further, our and previous (*22*) studies have shown that a subset of CL neurons maintains a high firing rate (40-50 Hz) during wakefulness. Mimicking this firing rate under anesthesia may partially explain the increased efficacy of gamma stimulation found in our data and another study (*23*). The frequency and location specificity of our stimulation effects suggest that clinical deep brain stimulation can be optimized to better reflect the desired neural dynamics of affected thalamocortical circuits to help alleviate disorders of consciousness.

## Acknowledgments

We thank M.I. Banks, C.W. Berridge, R.A. Pearce, R.D. Sanders, B.M. Krause, and C. Murphy for useful comments on the manuscript.

## Author contributions

M.R., J.P., N.K., S.M., A.R. and Y.S. performed research; M.R., S.A., G.D., M.A. and Y.S. analyzed data; M.R., A.R. and Y.S. wrote paper; M.R., J.P., N.K., S.M., S.A., G.D., M.A., A.R. and Y.S. edited paper.

## Data and materials availability

All data and code available upon reasonable request.

## Supplementary Materials

### Materials and Methods

#### Animals

The University of Wisconsin-Madison Institutional Animal Care and Use Committee approved all procedures, which conformed to the National Institutes of Health Guide for the Care and Use of Laboratory Animals. We acquired data from two male monkeys (*Macaca mulatta,* 4.3-5.5 years old, 7.63-10.30 kg body weight).

#### Neuroimaging

We performed structural imaging on anesthetized monkeys using the GE MR750 3T scanner (GE Healthcare, Waukesha WI). At the start of each scan session, we pre-medicated the monkey with ketamine (up to 20 mg/kg body weight) and atropine sulfate (0.03-0.06 mg/kg), prior to intubation. We then administered isoflurane (1-3% on ∼1 L/min O_2_ flow) to the monkey, with a semi-open breathing circuit and spontaneous respiration, to maintain general anesthesia for the duration of the session. We monitored the monkey’s vital signs (expired carbon dioxide, respiration rate, oxygen saturation, pulse rate, temperature) using an MR-compatible pulse oximeter and rectal thermometer.

We acquired a high-resolution structural brain image prior to the implant surgery, to delineate thalamocortical regions of interest (ROIs), and, after craniotomy, additional scans of electrodes *in situ* to confirm electrode positioning. For these three dimensional T1-weighted structural images, we used an inversion-recovery prepared gradient echo sequence with the following parameters: FOV=128 mm^2^; matrix=256 x 256; no. of slices=166; 0.5 mm isotropic; TR=9.68 ms; TE=4.192 ms; flip angle=12°; inversion time (TI)=450 ms). To generate the high-quality structural image, we collected 6-10 T1- weighted structural images and calculated the average image for each monkey using the FMRIB Software Library (FSL) (*24*). To localize electrodes, we averaged two structural images of electrodes *in situ*.

#### Surgery

We induced anesthesia with ketamine (up to 20 mg/kg body weight, i.m.) and maintained general anesthesia during aseptic surgical procedures with isoflurane (1-2%). We used 12 ceramic skull screws and dental acrylic to affix head implants on monkeys. We drilled 2.5 mm craniotomies in the frontal and parietal bones within a customized plastic recording chamber, providing access to our three thalamocortical ROIs in the right hemisphere: frontal eye field (FEF), lateral intraparietal area (LIP), and central lateral thalamic nucleus (CL). We derived craniotomy coordinates from the high-quality T1- weighted structural images acquired prior to the surgery. We fitted each craniotomy with a conical plastic guide tube filled with bone wax (guide tube prefabricated using model of skull based on T1-weighted structural images) (*25–27*) through which linear electrode arrays traversed. We also inserted two titanium skull screws within the recording chamber, one from which to record the EEG and one to serve as a reference. The head implant included a head post and, on the implant left and right sides, four hollow slots (two on each side) into which rods fitted, allowing head immobilization during electrophysiological recordings.

#### Behavioral tasks and sensory stimuli

To compare electrophysiological data between different states of consciousness, we needed to acquire data under similar behavioral and sensory conditions for awake and anesthetized monkeys. Thus, we acquired electrophysiological data from both awake and anesthetized monkeys during a passive auditory oddball paradigm as well as during “resting state” (in which no sensory stimuli were presented). The passive auditory oddball paradigm was useful because it does not require a behavioral response, does not require open eyes, and auditory stimuli have been shown to elicit neuronal responses from FEF (*28–30*) and LIP (*31–34*), allowing sound-aligned current source density analyses. Additionally, as controls in the awake monkeys, we acquired electrophysiology data during a fixation task, and during the passive auditory oddball paradigm while the monkey maintained fixation (oddball paradigm run concurrently with fixation task; see “Awake experiments” section). All electrophysiological recordings occurred in a quiet, dark room.

In the passive auditory oddball paradigm, the sequence of auditory tones (200 ms duration, with 800 ±100 ms jitter between each tone) comprised 80% standard tones (0.9 kHz frequency) and 20% deviant/oddball tones (1 kHz frequency). At least the first four stimuli of a sequence (3 min duration for anesthesia; 6 min duration for wake) were standard tones, and two sequential tones could not be deviant stimuli, otherwise the tone order was pseudorandom within the constraint of the overall 80/20 standard-to-deviant ratio. We presented tones using two speakers placed 35 cm from each ear under anesthesia and 80 cm from each ear during wakefulness (sound level at each ear was about 75 dB SPL for both states).

In the fixation task, the monkey needed to fixate a central fixation point (dim gray circle of diameter 0.42 degrees of visual angle on black background) on the monitor screen located 57 cm away. The monkey received a small volume (0.18-0.22 mL) of juice every 2.2-3.5 s while maintaining fixation within a 3 x 3 degree of visual angle window, centered on the fixation point. When the monkey’s gaze left the fixation window, he would typically re-establish central fixation quickly, to again receive juice every 2.2-3.5 s while fixating. To encourage long fixations, we doubled the juice volume if fixation persisted beyond 10 s. We only analyzed electrophysiological data during stable eye epochs (eye position remained fixed for at least 1 s). This applied to all wake-state data (resting, oddball paradigm and fixation task).

For awake experiments, we monitored monkeys’ eye position using a video-based eye tracker (500 Hz sampling rate). For anesthesia experiments, we monitored eyes using a digital video camera (capturing 30 frames per second) and used MATLAB to analyze luminance contrast in a window tightly bounding the eye image. The contrast differentiated closed eyes (i.e., relatively homogenous high luminance eyelid shade) and thalamic stimulation-induced eye openings (i.e., dark pupil and iris contrasting against sclera), as shown in Fig. 1A; and visual inspection of the eye video verified the timing of eye openings/closings derived from the contrast analysis.

#### Arousal scoring

We developed an arousal index based on clinical arousal scales to measure the behavioral effects of electrical stimulation. The arousal index incorporated five main indicators of arousal, with each indicator scored 0, 1 or 2, and the sum of the scores of the five indicators yielding the arousal index (range 0-10). The five indicators are:

1. limb/face movements (0 = nothing; 1 = small movement or increased EMG with no clear movement; 2 = full reach or withdrawal)
2. oral signs (0 = nothing; 1 = small mouth/jaw/tongue movements; 2 = full jaw openings/closures, with multiple repetitions)
3. body movements (0 = nothing; 1 = small torso movement or swallowing; 2 = large full torso movement)
4. eye movements/openings (0 = nothing; 1 = eyelid flutters/small blinks or increased eye movements; 2 = full eye opening with occasional blinks)
5. vital signs (0 = no change, i.e., difference of <10% respiration rate (RR), <5% heart rate (HR); 1 = difference of >10% RR, >5% HR; 2 = at least 20% change in either RR or HR, or at least 10% change in both RR and HR; compared to baseline 30 s prior to stimulation).

A veterinarian at the Wisconsin National Primate Research Center, a clinical anesthesiologist, and five other primate electrophysiologists observed the electrical stimulation effects during anesthesia experiments. Using observations recorded at the time of stimulation experiments as well as offline review of videos and EMG data (filtered 30-450 Hz, full-wave rectified, then filtered 5-100 Hz to extract the envelope), we scored arousal level prior to, during, and after all stimulation events. A typical stimulation block consisted of three stimulation event repetitions (one minute each) within a seven minute recording period at a given site, using the same stimulation frequency, current, polarity, duration, anesthetic and dose. We defined stimulation event epochs from the onset to offset of pulses, i.e., from 1-2 minutes, 3-4 minutes, and 5-6 minutes of a seven minute block. The time between two stimulation epochs was split equally into post- and pre-stimulation epochs (see Fig. 1A for an example). The pre-stimulation, during stimulation and post-stimulation arousal index for a block reflected the maximum possible score across the repetitions (repetitions largely produced the same score within each epoch type). Prior to electrical stimulations (except for a rare few instances testing the valence of different stimulation frequencies), the arousal index was 0 or 1. This could be differentiated from stimulation events inducing an arousal index of 3 or more by all observers. Thus, we defined effective stimulation events as those inducing an arousal index of 3 or more, whereas ineffective stimulation events had an arousal index of 0-2.

#### Electrophysiological recording and electrical stimulation details

FEF and LIP electrodes had either 16 or 24 contacts, and CL electrodes had 24 contacts. These platinum/iridium electrode contacts had a diameter of 12.5 μm, and 200 μm spacing between contacts. The impedance of contacts on recording electrodes was typically 0.8-1 MΩ. We also measured the EEG using titanium skull screws located above dorsal fronto-parietal cortex and, in anesthetized experiments, the EMG using a hypodermic needle (30G) in the forearm. We recorded electrode signals (filtered 0.1- 7,500 Hz, amplified and sampled at 40 kHz) using a preamplifier with a high input impedance headstage and OmniPlex data acquisition system controlled by PlexControl software.

We electrically stimulated using 24-contact electrode arrays that had previously been used several times as recording electrodes (and now had lower impedance). In early stimulation trials, we titrated current (tested 100-300 μA, but because 100-200 μA induced arousal, there were only a small number of >200 μA cases), polarity of first phase of biphasic pulse (negative- or positive-going first phase), number of electrode contacts simultaneously stimulated (tested 1, 4, 8 and 16 contacts), and stimulation duration (15-60 s). For subsequent electrical stimulations, we simultaneously stimulated via 16 electrode contacts, with 400 μs bi-phasic pulses of 200 μA, for a total of 60 s stimulation duration for any given stimulation event (experiments included multiple stimulation events). We typically performed three stimulation events at a given frequency within a stimulation block for reproducibility, with a recovery time of at least the stimulation event duration between repetitions, i.e., stimulations from 1-2 minutes, 3-4 minutes, and 5-6 minutes of a seven minute block. In our analyses, we included all stimulation data with currents from 100-200 μA. Stimulation event duration, ranging from 15-60 s, did not influence arousal indices, so we included all durations in our analyses.

#### Electrode array localization

We acquired T1-weighted structural images with electrodes held *in situ* by the customized guide tubes (*27*). While the actual electrode is not visible in the images, a susceptibility “shadow” artifact appears along the length of the electrode with a width of approximately one voxel (0.5 mm^3^, either side of the electrode). We targeted electrodes to thalamocortical ROIs based on the individual monkey’s structural images, using a stereotaxic atlas as a general reference (*35*). We re-positioned electrodes as necessary and re-acquired T1-weighted structural scans until electrodes were in their desired locations in the thalamus and cortex. Offline, we registered (6 degrees of freedom) the images with electrodes *in situ* to the high-quality structural image acquired prior to surgery. Using measurements of electrode depth during imaging and recording sessions as well as the image of electrodes *in situ*, we reconstructed recording and stimulation sites along electrode tracks. Thalamic stimulation sites, specifically the eighth electrode contact of the 16 contacts simultaneously used for electrical stimulation (i.e., middle of stimulating array), are shown on one coronal slice (sites collapsed across the anterior-posterior axis) in Fig. 1 (monkey R).

We further validated the localization of recording sites in our three thalamocortical ROIs using functional criteria. We confirmed the FEF ROI in an initial experiment using electrical stimulation at the frontal recording site, i.e., low currents (<100 μA) elicited eye movements (36). In the LIP ROI during awake experiments, a large number of neurons showed the classical response characteristic of peri-saccadic activity. In the CL ROI, we found a subset of neurons with high firing rates (around 40-50 Hz) in the awake state, consistent with a CL locus (22).

With the aim of positioning electrode contacts in all cortical layers in FEF and LIP, we used depth measurements derived from structural images to initially position electrode arrays across FEF and LIP layers (24 contacts with 200 μm spacing between contacts corresponds to a 4.6 mm span, and 16 contacts correspond to a 3 mm span, which generally allows for contacts in superficial, middle and deep cortical layers for tracks near perpendicular to the cortical surface or with moderate angles from perpendicular). We further adjusted electrode position to maximize the number of contacts showing single-unit or multi-unit spiking activity, and we visualized evoked potentials to auditory tones, with middle layers showing earliest response. We then used current source density (CSD) analysis to attribute contacts to superficial, middle and deep cortical layers (see section on CSD analysis below).

We performed post-mortem histology to reconstruct electrode tracks in one monkey (in addition to the reconstructions using structural MRI and electrode depth measurements in both monkeys). After fixing the brain in 10% neutral buffered formalin, the right hemisphere was cut into approximately 5 mm thick coronal sections, embedded in paraffin, then thinly sectioned (8 μm). Around ROIs, we stained sections with Hematoxylin and Eosin, and visualized sections under a microscope to confirm electrode tracks through our ROIs.

We recorded 282 CL neurons, 281 FEF neurons and 282 LIP neurons in total. For CL, there were 181 neurons during anesthesia; 101 neurons during wakefulness; and 83 neurons during sleep. For FEF superficial, middle and deep layers, there were respectively 48, 33 and 91 neurons during anesthesia; 37, 22 and 50 neurons during wakefulness; and 37, 22 and 42 neurons during sleep. For LIP superficial, middle and deep layers, there were respectively 38, 34 and 91 neurons during anesthesia; 36, 10 and 73 neurons during wakefulness; and 24, 9 and 65 neurons during sleep. Neurons recorded during sleep were also recorded during the wake state. Neurons recorded during anesthesia were recorded in different sessions from neurons recorded during wakefulness/sleep.

#### Anesthesia experiments

We used either isoflurane (9 sessions: 5 for Monkey R, 4 for Monkey W) or propofol (9 sessions: 4 for Monkey R, 5 for Monkey W) in anesthesia experiments, to ensure that results were not drug-specific, instead reflecting general mechanisms of anesthesia/consciousness. The duration of each anesthesia experimental session was 10-12 hours. We induced anesthesia with ketamine (up to 20 mg/kg body weight, i.m.), then intubated the monkey and inserted an intravenous catheter(s) for fluid and drug administration. We maintained general anesthesia in spontaneously respiring monkeys with isoflurane (0.8-1.5% on 1 L/min O_2_ flow) or propofol (0.17-0.33 mg/kg/min i.v.), and a clinical anesthesiologist (A.R.) oversaw stable conditions throughout. We categorized doses as lower (isoflurane < 1%; propofol < 0.23 mg/kg/min), medium (isoflurane 1- 1.19%; propofol 0.23-0.26 mg/kg/min) and higher (isoflurane ≥ 1.2%; propofol ≥ 0.27 mg/kg/min) within the aforementioned ranges for statistical purposes (see “Statistical analysis” section). We positioned monkeys in the prone position within a modified stereotaxic apparatus atop a surgical table, with the monkey’s head immobilized by four rods (attached to the stereotaxic device) that slid into the implant hollows. We maintained the monkey’s temperature using a forced-air warming system and monitored vitals (end tidal carbon dioxide, respiration rate, oxygen saturation, heart rate, blood pressure and rectal temperature).

Each experimental session had two parts: the first part involved simultaneous recordings from FEF, LIP and CL (recordings started at least two hours after anesthetic induction and ketamine administration), and the second part involved electrical stimulation of CL during simultaneous recordings from FEF and LIP without changing the anesthetic regimen. We independently positioned linear multielectrode arrays in each ROI, and allowed arrays to settle for 30 minutes prior to starting recordings. Microdrives coupled to an adapter system allowed different approach angles for each ROI. For both parts of experiments, we interleaved resting state epochs and the passive auditory oddball paradigm. During the first part of the experimental session, we performed neural recordings at a number of different anesthetic levels, adapting the dose to reflect a range of clinically relevant anesthetic depths, e.g., 1%, 1.1%, 1.25% and/or 1.5% isoflurane, or 0.2, 0.225, 0.25 and/or 0.3 mg/kg/min propofol, allowing dosing changes to stabilize before starting the next block of recordings (typically at least 30 minutes). During the second part of the experiment, we either electrically stimulated using the linear multielectrode array existing in the thalamus or replaced it with another array inserted along the same trajectory to the same depth. We first stimulated thalamic sites at a frequency of 50 Hz. If this did not induce arousal, then we moved the stimulating electrode to a new depth in the thalamus in steps of 0.5-1 mm dorsal or ventral along the electrode track, until stimulation induced arousal. When 50 Hz stimulation induced arousal, we tested additional stimulation frequencies, i.e., 2, 10 or 200 Hz, or further depths (mapping the area of effect). The order of stimulation frequencies generally followed one of two patterns: 50 Hz alternating with one of the other stimulation frequencies; or multiple repetitions of a particular stimulation frequency, followed by multiple repetitions of a different stimulation frequency.

In early experiments, we tested thalamic stimulations at different anesthetic doses between 0.8-1.3% for isoflurane and between 0.17-0.3 mg/kg/min for propofol. We observed thalamic stimulation-induced arousal for all but the highest doses (i.e., 1.3% isoflurane and 0.3 mg/kg/min propofol). In subsequent isoflurane experiments, we used doses between 0.8 – 1.25% (M = 1.04, SD = 0.11) during thalamic stimulation, and in propofol experiments, we used doses between 0.17-0.28 mg/kg/min (M = 0.23, SD = 0.03). Data for all doses were included in analyses and controlled for statistically (see “Statistical analyses” section).

As an additional control, we also separately stimulated the FEF and LIP using the same stimulation parameters as that used in the thalamus (10 or 50 Hz). FEF or LIP stimulation alone did not induce arousal.

#### Awake experiments

We performed 40 awake experimental sessions (18 for monkey R; 22 for monkey W), each session usually of 2-4 hours duration. Monkeys sat upright in a primate chair with their head immobilized using the head post and/or four rods that slid into the hollow slots in the head implant. Awake experiments were split into two types (similar to the two parts of anesthesia experimental sessions); those with and without thalamic stimulation. Experiments without stimulation involved simultaneous recordings from FEF, LIP and CL across multiple blocks of all task conditions. Stimulation experimental sessions involved electrical stimulation of CL, at different frequencies, during simultaneous recordings from FEF and LIP across all task conditions. During each type of experiment, we interleaved task conditions involving reward (fixation and oddball fixation) with those not involving rewards (resting state and passive oddball). The specific task order was varied randomly across different experimental sessions.

For electrical stimulation, we pseudorandomly applied stimulation blocks of different frequencies, i.e., 10, 50 and 200 Hz. Because electrical stimulation of the thalamus at 50 Hz frequency (or other frequencies) in awake monkeys did not elicit any movements (as observed during effective stimulation events in anesthesia experiments), it is unlikely that the effects of 50 Hz stimulation of CL in anesthetized monkeys simply reflected direct effects on the motor system. Rather, it supports the finding that 50 Hz stimulation effects reflected increased arousal.

We performed neural recordings from brain areas implicated in awareness. Because these areas are also involved in selective attention and oculomotor function, we aimed to ensure differences between wake and anesthesia results were not related to attentional or saccadic processes. To this end, within the wake state, we compared recordings during the fixation task to resting state, as well as recordings during the passive oddball with fixation to that without fixation. For each condition (fixation task, resting state, oddball with and without fixation), we analyzed epochs (at least 1 s in duration) in which the monkey’s eye position was stable, as verified using the eye tracker. These analyses showed neural data from compared conditions to be qualitatively similar. Considering these controls, to keep wake and anesthesia conditions as similar as possible, we compared wake and anesthesia data collected during conditions in which there were no task demands, i.e., the resting state and passive oddball conditions (not the fixation task or the oddball with fixation) in the dark.

#### Sleep

During awake experiments, monkeys at times would fall asleep, particularly during conditions not involving rewards, such as the resting state. Online, we identified non-rapid eye movement (NREM) sleep using the following criteria: increased delta (1-4 Hz) activity in EEG (compared with wake); extended eye closure (recording times when eyes closed and re-opened, to compare with semi-automatic detection offline); preceding period of drowsiness indicated by slow drooping/closing of eyelids; stop in fixation task performance (if current task is fixation task); and no overt body movement. Offline, we identified NREM sleep periods using EEG and eye tracker data. We bandpass filtered (1- 4 Hz; Butterworth, order 6) EEG data and applied the Hilbert transform, to calculate the instantaneous delta-band amplitude. From the resulting time series, we detected times of relatively high delta amplitude using thresholds titrated for each recording session, because the mean delta amplitude and standard deviation could vary depending on the recording session and total sleep time. For each session, we selected the threshold as the number of standard deviations from the mean delta amplitude that produced a total sleep time estimate that closely resembled the expected sleep time based on online NREM identification, as well as the offline calculation of the total time when the monkey’s eyes were closed (using the recorded eye tracker time series data). Offline NREM sleep identification and time stamping then involved automated detection of extended epochs across the recording session when both the monkey’s eyes were closed and delta amplitude was above threshold. These offline NREM sleep detections were similar to manual online detections, and proved reliable for different recording sessions and monkeys.

The identified sleep epochs corresponded to early phases of NREM sleep (N1 or N2, i.e., light sleep). Thus, monkeys were not at the same depth of unconsciousness during sleep as they were during general anesthesia in our study. This notwithstanding, we included the spike rate data during early NREM sleep, as this allowed us to compare the influence of conscious and less-conscious states on the same subset of neurons (n = 282) recorded in both wakefulness and sleep. This further substantiated our comparison of spiking activity between the awake and anesthetized states, activity recorded from two different samples of neurons from the same ROIs (maintenance of stable anesthesia up to 12 hours required recordings to take place in a surgical suite, whereas awake recordings took place in the behavioral lab). Because local field potentials (LFPs) reflect combined activity from a considerably larger volume (compared with single-neuron activity) (*37*), LFPs recorded at different times, i.e., awake and during anesthesia, are more readily compared. Nonetheless, we include early NREM sleep LFP data as well, to further substantiate the altered connectivity during anesthesia (although a complete account of sleep influence on our thalamocortical recordings is beyond the scope of this study).

#### Neural data preprocessing

We defined data segments of 1 s duration (akin to trials) for analysis. In the awake state, we first determined stable eye epochs (to match eye behavior between conscious and unconscious states), i.e., epochs starting 200 ms after a saccade and ending 200 ms before the next saccade. Next, we divided stable eye epochs into non-overlapping 1 s windows. In the anesthetized and non-REM sleep states (when eyes are closed), we divided all data in each of these states into non-overlapping 1 s windows.

We lowpass filtered data to 250 Hz for LFPs (Butterworth, order 6, zero-phase filter). Next, we linearly detrended LFPs, then extracted artifacts from LFP data, by removing significant sine waves using the Chronux function rmlinesc. Individual electrode contacts with signal amplitude greater than 5 standard deviations from the mean were excluded from analysis. For power and coherence analyses, we further calculated bipolar derivations of LFPs, i.e., the difference between two adjacent electrode contacts (excluding contacts that had been removed due to noise), to minimize any possible effects of a common reference and volume conduction (*38–40*).

We bandpass filtered data 250-5,000 Hz for spiking activity (Butterworth, order 4, zero-phase filter) and sorted spikes using Plexon Offline Sorter software. Initial spike detection involved thresholding data at >3 standard deviations away from the mean. We then used principal components analysis to extract features of the spike shapes. Finally, we used the T-distribution expectation maximization algorithm to identify clusters of spikes with similar features.

For neural data during electrical stimulation, there was a brief artifact caused by the applied current. To remove this artifact, we first excised a 1 ms window around the artifact, then linearly interpolated across this window. Next, we used the Chronux function rmlinesc to remove any significant sine waves at the stimulation frequency (we also performed artifact removal using the SARGE toolbox (*41*), which yielded qualitatively similar results).

#### Spike rate

We calculated the average spike rate in 1 s windows (during stable eye epochs) for each neuron, in the awake, sleep and/or anesthetized states. We divided anesthetized state data into electrical stimulation and no stimulation windows. For electrical stimulation data, we calculated the spike rate during the stretches of data unaffected by the stimulation-induced artifact.

#### Spike timing

For each neuron, we generated interspike interval (ISI) histograms (1 ms bin width), from which we derived an index of burst firing propensity in the awake and anesthetized states (*42*). We excluded neurons with very low spike rate (< 1 Hz) from the burst index analysis, as their ISI histograms had too few samples). For thalamic neurons, the burst index equaled the proportion of spikes occurring within 2-8 ms (sum of spikes in the 2-8 ms bins of the ISI histogram divided by the total number of spikes; Fig. 2D); we also calculated indices for 2-5, 2-10 and 2-15 ms bins (for qualitatively similar results). Because ISIs in CL neuronal bursts have been reported to commonly range up to 6 ms (lengthening with increasing burst size) (*43*), we selected the next accommodating window size, 2-8 ms. For cortical neurons, the burst index equaled the proportion of spikes occurring within 2- 15 ms (sum of spikes in the 2-15 ms bins of the ISI histogram divided by the total number of spikes; Fig. 2E); we also calculated indices for 2-10, 2-20 and 2-30 ms bins. Although it has been reported that LIP neurons have a low tendency to burst in the wake state (*44*), we still measured changes in spiking regularity across different states, by using a relatively larger window, 2-15 ms (cf. CL), still applicable for frontal cortex (45), to allow comparisons between cortical areas.

#### Current source density (CSD)

We localized electrode contacts to superficial, middle or deep cortical layers based on inverse CSD analyses (46). To do this, we used the CSDplotter toolbox for MATLAB (https://github.com/espenhgn/CSDplotter; dt = 1 ms, cortical conductivity value = 0.4 S/m, diameter = 0.5 mm) for calculating the inverse CSD in response to auditory tones in the passive oddball paradigm. Linear multi-electrode arrays measure the LFP, *ϕ*, at N different cortical depths/electrode contacts along the z-axis with spacing h. The standard CSD, C_st_, is estimated from the LFPs using the second spatial derivative, i.e.,

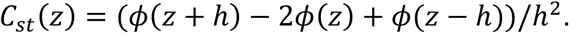

LFPs can also be estimated from given CSDs, represented in matrix form as Φ=F*Ĉ*, where Φ is the vector containing the N measurements of ϕ, *Ĉ* is the vector containing the estimated CSDs, and F is an NxN matrix derived from the electrostatic forward calculation of LFPs from known current sources. The inverse CSD method uses the inverse of F to estimate the CSD, i.e., *Ĉ*=F^-1^ Φ. For the step inverse CSD method (*46*) used here, it is assumed that the CSD is step-wise constant between electrode contacts, so the sources are extended cylindrical boxes with radius R and height h. In this case, F is given by:

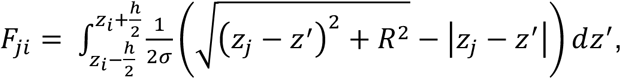

where σ is the electrical conductivity tensor, and ϕ(z_j_) is the potential measured at position z_j_ at the cylinder center axis due to a cylindrical current box with CSD, C_i_, around the electrode position z_i_. The inverse CSD method offers advantages over the standard CSD. The inverse CSD method estimates the CSD around all N electrode contacts, whereas the standard CSD method yields estimates around N-2 contacts. Further, the standard CSD requires equidistant contacts, whereas the inverse CSD method does not, which is advantageous when data from a noisy contact may need to be excluded. We used the step inverse CSD method, because it may perform better than the delta-source CSD method as electrode contact spacing increases, and the spline CSD method can be more sensitive to spatial noise, e.g., from gain differences between electrode contacts or from an excluded contact (*46*).

We identified the early current sink in response to auditory stimulation and designated the bottom of the sink as the bottom of the middle layers (around boundary between layers 4 and 5). We included the electrode contact at the bottom of the middle layers and the two more superficial contacts as the middle layers. Electrode contacts in FEF or LIP superficial to the middle layers were designated as being in the superficial layers, whereas FEF or LIP contacts deeper than the middle layers were designated as being in the deep layers. Layer assignments were cross-referenced to reconstructions of the recording sites along the electrode track (based on measurements of electrode depth as well as the image of electrodes *in situ*) as well as to single-unit or multi-unit spiking activity, which helped delineate the border between gray and white matter. We excluded from analysis contacts that were found to be located outside the ROI.

Previous studies generated CSD data in FEF (*47*) and LIP (*48*) using visual stimulation (FEF: 1 degree of visual angle square at 60% contrast; LIP: diffuse light). Our CSD profiles generated with auditory stimulation were roughly consistent with these previous studies of FEF and LIP in so far as sensory stimulation elicited early sinks in middle layers (which would be predicted based on auditory stimulation activating middle cortical layers relatively early).

We also performed CSD analyses, in the case of resting state recordings, using LFP signals aligned to the trough of delta-band oscillations recorded from the electrode contact with the highest delta power (i.e., this contact served as the phase index) (*8, 39, 40*). These delta phase-realigned CSDs showed differences across cortical layers which helped verify that probe positions remained stable across recording blocks that did not include auditory stimuli of the passive oddball paradigm.

#### Power

We calculated power in 1 s windows (stable eye epochs) for every bipolar-derived LFP, using multi-taper methods (5 Slepian taper functions, time bandwidth product of 3, averaging over windows/trials) with the Chronux data analysis toolbox for MATLAB (http://chonux.org/) (*49, 50*). Noisy trials, samples with amplitudes that exceeded 4 standard deviations from the mean, were removed. Sinusoidal noise, especially at stimulation frequencies and 60 Hz, was removed using notch filters or the Chronux function rmlinesc. There were an unequal number of windows per condition, due to differences in data length and number of stable eye epochs. Because the number of time windows (or trials) affects the power estimate (S(f)), we bias-corrected power values (*58*). The bias-corrected power spectrum, B(f), is given by:

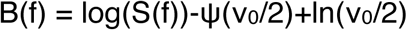

where ν_0_ = 2*K*N, where K is the number of tapers (5) and N is the number of time windows. To obtain population values, we pooled the bias-corrected power estimates for the awake state and again for the anesthetized state (separately for the no stimulation, effective stimulation and ineffective stimulation conditions).

#### Coherence

We calculated coherence using multi-taper methods (5 Slepian taper functions, time bandwidth product of 3) with the Chronux toolbox. Noisy trials, samples with amplitudes that exceeded 4 standard deviations from the mean, were removed. Sinusoidal noise, especially at stimulation frequencies and 60 Hz, was removed using notch filters or the Chronux function rmlinesc. We used the coherence measure to study the temporal relationship between LFPs, or between spikes and LFPs, within and between the thalamus, FEF and LIP. The coherence is given by:

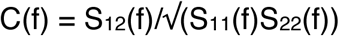

where S(f) is the spectrum with subscripts 1 and 2 referring to the simultaneously recorded spike/LFP at one site and LFP at another site. The coherence is normalized between 0 and 1, so it can be averaged across different pairs of time series. For each paired recording, we calculated the coherence in 1 s windows during which the monkey’s eyes were stable. There were an unequal number of windows per condition, due to differences in data length and number of stable eye epochs. Because the number of time windows (or trials) affects the coherence estimate, we bias-corrected/transformed coherence values (*51*). The transformed coherence, T(f), is given by:

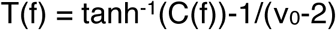

where ν_0_ is the degrees of freedom; for our multi-taper estimates, ν0 = 2*K*N, where K is the number of tapers (5) and N is the number of time windows. To obtain population values, we pooled the transformed coherence estimates for the awake state and again for the anesthetized state (separately for the no stimulation, effective stimulation and ineffective stimulation conditions).

To ensure that changes in coherence did not simply reflect changes in power at given frequency bands, we investigated the relationship between our power and coherence results. While we did find that anesthesia increased delta power and decreased power at higher frequencies for all cortical areas, power changes during thalamic stimulation were broadband and typically smaller for effective relative to ineffective stimulations (unlike coherence changes; Fig. S6; Table S3 and S4). This poor correlation between arousal and power during stimulation suggests that power is unlikely to be driving stimulation-induced changes in coherence, and is not a key component of the NCC.

#### Statistical analyses

We performed statistical analyses using general linear models (GLMs) in R, regressing the relevant dependent variable on all independent variables, interactions, and covariates (models M1-M21 below). We used linear models (LM in R) for effects that varied between all other effects, yielding T statistics for each estimated slope (β parameter). Effects that varied within other effects of interest were estimated using linear mixed effect models (LMER in R), yielding F statistics, or after computing difference scores with linear models, yielding T statistics. Random effects of LMER models are represented as gamma parameters, and all simple effects are presented as beta parameters, where the slope for the effect of interest is β_1_. P values stemming from the same family of statistical tests (models intended to describe the same effect in different populations) were controlled for multiple comparisons using Holm’s correction.

To compare doses between anesthetics, we separated doses into lower (−1), medium (0), and higher (1) dose groups within the experimental range used for both anesthetic agents. For isoflurane, lower doses were < 1%, medium were ≥ 1% and < 1.2%, and higher doses were ≥ 1.2%. For propofol, lower doses were < 0.23 mg/kg/min, medium were ≥ 0.23 and < 0.27 mg/kg/min, and higher doses were ≥ 0.27 mg/kg/min. This allowed us to use coded dose (DoseCode) as a covariate independent of anesthetic.

To contrast stimulation effectiveness, we coded stimulations producing arousal ≥ 3 as effective (1) and those producing arousal < 3 as ineffective (0). This allowed us to compare neural dynamics across stimulations that reflected clear changes in the level of consciousness while controlling for changes that may be induced only by introduction of thalamic current, which was the same for ineffective and effective stimulations.

To limit the number of multiple comparisons across frequency, we averaged power and coherence across canonical frequency bands: delta = 0-4 Hz, theta = 4-8 Hz, alpha = 8- 15 Hz, beta = 15-30 Hz, low gamma = 30-60 Hz and high gamma = 60-90 Hz. As a control for possible artifacts, we also averaged more selectively within the low gamma (across 30-47 and 53–57 Hz) and high gamma (63-90 Hz) bands, so as not to include data at 50 Hz, the frequency of thalamic stimulation, and 60 Hz, the frequency of power line noise, producing similar results.

##### Stimulation effects

To test the general effect of thalamic stimulation on arousal (Fig. 1B), we regressed arousal score within stimulation blocks on the peri-stimulation epoch (pre, stimulation, post), including dose and anesthetic as covariates. Peri-stimulation epoch (StimEpochF) was dummy coded as a factor referenced to the epoch with stimulation, anesthetic was coded as a centered dichotomous variable (isoflurane = −0.5, propofol = 0.5), and dose was treated as DoseCode. We included random slopes only for stimulation epochs, as dose and anesthetic remained constant within a given stimulation event. Significantly negative β_1_ shows that outside of stimulation, arousal score is lower even controlling for the effects of dose and anesthetic.

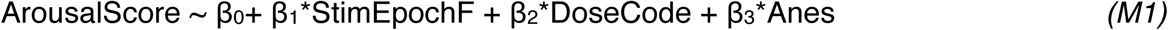

To ensure the effects of stimulation were not being driven or modulated by dose, we repeated this model using DoseCode as an interactant, but the interaction was not significant (Fig S1., A-C).

To test the effect of dorsal-ventral (D-V) proximity to CL of the stimulation array on arousal (Fig. 1F), we regressed the stimulation arousal difference (stim – pre) on the linear and quadratic components of the D-V proximity to CL of the centermost contact of each stimulation array, including dose, anesthetic, and variation of placement along the medial-lateral (M-L) axis as covariates. D-V distance (D-Vdist) was coded as the distance from the centermost contact of the stimulation array to center of CL in the D-V plane in each monkey, dose was coded as DoseCode, anesthetic (Anes) was coded as a dichotomous variable (isoflurane = −0.5, propofol = 0.5), and M-L distance (M-Ldist) was coded as the linear distance from the centermost contact in the stimulation array to the center of CL in the M-L plane for each animal. Significantly negative β_1_ shows that moving more dorsal or more ventral from the center of CL decreases the arousal induced by stimulation above and beyond the effects of anesthetic, dose, and M-L variation.

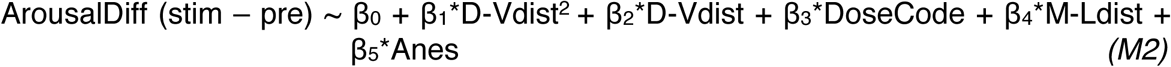

To test the effect of Euclidian distance from the center of CL on arousal (Fig. S1, G-I), we regressed arousal difference (stim – pre) on the Euclidian distance (Distance) including dose, anesthetic, and monkey as covariates. Importantly, we included monkey as a covariate in this model to control for general differences between the monkeys in terms of the size and shape of their anatomy in M-L and D-V planes (main source of variation in our electrode track locations). Euclidian distance was calculated as the length of a vector from the center of CL to the centermost contact of each stimulation array. Dose was coded as DoseCode, anesthetic (Anes) was coded as a centered, dichotomous variable (isoflurane = −0.5, propofol = 0.5), and monkey (Animal) was coded as a centered, dichotomous variable (monkey R = −0.5, monkey W = 0.5). Significantly negative β_1_ shows that moving further from the center of CL in any M-L/D-V direction decreases the arousal induced by stimulation above and beyond variation contributed by dose, anesthetic, or monkey.

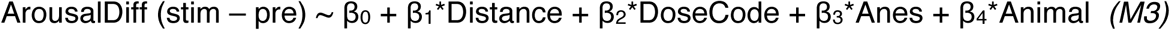

To test the relative effectiveness of stimulation frequency on arousal (Fig. 1H), we regressed arousal difference (stim – pre) on stimulation frequency, for all case-matched examples in which stimulations occurred at the same site, at multiple frequencies, and at least one stimulation had been effective (arousal score ≥ 3). Because only 50 Hz stimulations reliably increased arousal (Fig. 1H; error bars did not include 0), we coded stimulation frequency (StimFreq) as a dichotomous variable, where stimulations were either at 50 Hz (0.5) or not at 50 Hz (−0.5). Dose (DoseCode) was coded as a factor reflecting lower, medium and higher doses within our experimental range, and anesthetic (Anes) was coded as a centered dichotomous variable (isoflurane = −0.5, propofol = 0.5). Significantly positive β_1_ shows that, as monkeys go from pre to stimulation conditions, arousal increases more when stimulations are at 50 Hz above all other frequencies, even controlling for differences in dose and anesthetic between stimulation blocks.

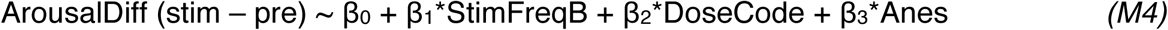

##### Spike rate effects

For non-stimulation data, we limited all comparisons to resting state and anesthesia conditions without auditory stimuli. To test the effect of sleep on thalamic spike rate (Fig. 2C), we regressed spike rate within neuron on state (wake vs sleep). State was coded as a dichotomous variable (wake = 0, sleep = 1). A random intercept and slope for state was included by neuron. Significant negative β_1_ shows that after neurons transition from wake to sleep, spike rate tends to decrease.

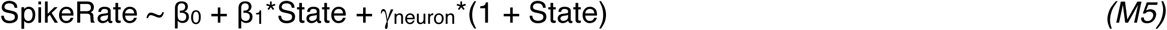

In the cortex (Fig. 2, F-H), we used a similar model but included the layer from which the neuron was recorded (SpikeLayer) as an interaction. SpikeLayer was dummy coded as a factor referenced to the deep cortical layers. A random intercept and slope for state was included by neuron. No random slope was included for spike layer as it could not vary within neuron. Finding significant positive β_1_ for the interaction of state and layer shows that the decrease in spike rate predicted by the state change is less for superficial relative to deep cortical layers controlling for variation in the middle layer spike rate. This model was used separately for neurons found in FEF and LIP (and controlled for multiple comparisons).

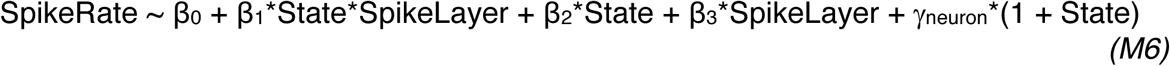

To test the effect of anesthesia on thalamic spike rate (Fig. 2C), we regressed spike rate between neuron on state (wake vs anesthesia). State was coded as a dichotomous variable (wake = 0, anesthesia = 1). Significant β_1_ shows that neurons recorded during anesthesia had lower spike rates relative to wakefulness.

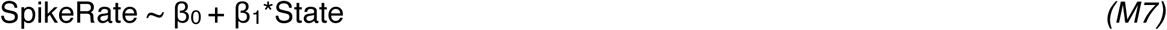

In the cortex (Fig. 2, F-H), we used a similar model but included the layer from which the neuron was recorded (SpikeLayer) as an interaction. SpikeLayer was dummy coded as a factor referenced to the deep cortical layers. Finding significant positive β_1_ for the interaction of state and layer shows that the decrease in spike rate predicted by the state change is less for superficial relative to deep cortical layers controlling for variation in the middle layer spike rate. This model was used separately for units found in FEF and LIP.

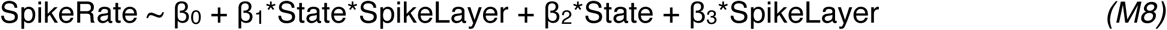

To ensure that effects were consistent between anesthetics, we compared the spike rate for each type of neuron (superficial, middle, deep cortical, or thalamic) separately across anesthetic states (isoflurane vs propofol; Fig. S2) including dose and cortical area as covariates. State was dummy coded as a dichotomous variable (propofol = 0, isoflurane = 1). Cortical area (Area) was coded as a centered dichotomous variable (FEF = −0.5, LIP = 0.5) and dose was coded as DoseCode. Negative β_1_ shows that spikes recorded during isoflurane have lower spike rate than those recorded under propofol (though none were significant after controlling for multiple comparisons).

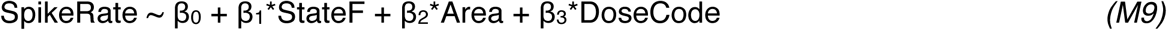

For stimulation data, we analyzed passive auditory oddball paradigm data in addition to resting state data. To test the effect of thalamic stimulation (50 Hz) on cortical spike rate (Fig. 1, I-K), we regressed spike rate within stimulation block on the 4-way interaction between peri-stimulation epoch (pre vs stim), cortical layer (superficial vs deep), stimulation effect (effective vs ineffective), and cortical area (FEF vs LIP), including dose, anesthetic and task as covariates. Peri-stimulation epoch (StimEpoch) was coded as a dichotomous variable (pre = −1, stim = 0), spike layer (SpikeLayer) was coded as a dichotomous variable (superficial = 0, deep = 1), stimulation effectiveness (StimEffect) was coded as a dichotomous variable (ineffective = 0, effective =1), and cortical area (Area) was coded as a centered dichotomous variable (FEF = −0.5, LIP = 0.5). In addition, dose was coded as DoseCode, and anesthetic (isoflurane = −0.5, propofol = 0.5) and task (resting state = −0.5, passive oddball = 0.5) were coded as a centered, dichotomous variables. A random intercept and slope for stimulation epoch was included by stimulation block (stimID), as this was the only variable which changed within a given stimulation block. A significantly positive β_1_*for the 4-way interaction shows that effective stimulation increases spike rate more for deep layers in LIP than any other condition, controlling for differences in dose, anesthetic, and task conditions.

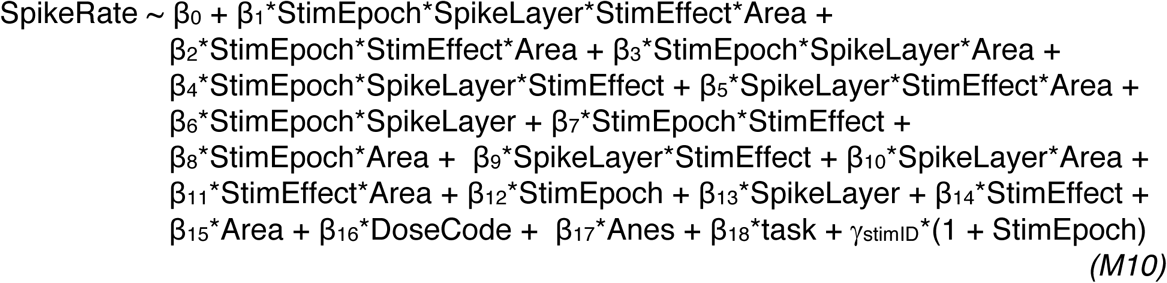

##### Bursting effects

To test the effect of sleep on thalamic bursting (Fig. 2D), we regressed bursting index within neuron (BI_2-8_; derived from the 2-8 ms bins of the ISI histogram) on state (wake vs sleep), including spike rate as a covariate (as spike rate tended to change with state in thalamic neurons and could influence the burst index). State was coded as a dichotomous variable (wake = −0.5, sleep = 0.5). Because the relationship between bursting changes and spike rate was largely linear within neuron, spike rate was coded as a continuous variable (total spikes/total time). We included a random slope for state and spike rate by neuron. A significant positive β_1_ indicates that after transitions from wakefulness into NREM sleep, thalamic neurons increase bursting, controlling for changes in spike rate.

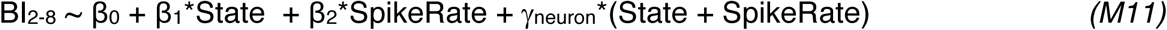

To test the effect of anesthesia on thalamic bursting (Fig. 2D), we regressed bursting index (BI_2-8_; derived from the 2-8 ms bins of the ISI histogram) between neuron on state (wake vs anesthesia), including spike rate as a covariate (as spike rate tended to change with state in thalamic cells and could influence the burst index). State was coded as a dichotomous variable (wake = 0, anesthesia = 1). As the relationship between bursting changes and spike rate were not reliably linear between neurons, spike rate was log transformed (SpikeRateL) and coded as a continuous variable (ln(total spikes/total time)). A significant positive β_1_ indicates that neurons recorded during anesthesia have higher burst index than wakefulness, controlling for differences in spike rate.

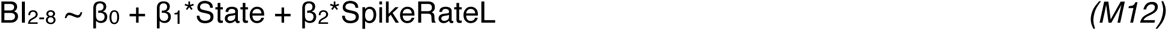

To test the effect of anesthesia on cortical bursting (Fig. 2E), we regressed bursting index (BI_2-15_; derived from the 2-15 ms bins of the ISI histogram) between neuron on the interaction of state (wake vs anesthesia) and spike layer, including spike rate and cortical brain area as covariates. State (wake = 0, anesthesia = 1) and spike layer (superficial = 0, deep = 1) were coded as dichotomous variables. As both cortical areas yielded similar results, we combined data across the cortex, and included cortical area as a centered, dichotomous covariate (FEF = −0.5, LIP = 0.5). Because the relationship between bursting changes and spike rate were not reliably linear between cells, spike rate was log transformed (SpikeRateL) and coded as a continuous variable (ln(total spikes/total time)). Significant positive β_1_ for the state and layer interaction indicates that the increased bursting during anesthesia is larger for deep relative to superficial neurons, controlling for differences in spike rate and cortical area.

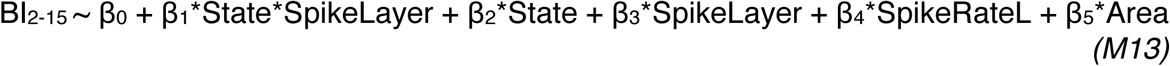

##### Power and coherence effects

We limited all non-stimulation comparisons to resting state and anesthesia conditions without auditory stimuli. To test the effects of anesthetics on power and coherence (Fig. 3, Fig. 4, Fig. S3, Fig. S4, Fig. S5, Fig. S6 and Fig. S7), we regressed power (S), coherence (C), and spike-field coherence (spikeFC) averaged across different frequency bands (delta, alpha, theta, beta, low gamma and high gamma) and isolated to different electrode contact pairs of interest (in the case of coherence, e.g., isolated to superficial-deep contact pairs within a cortical area, or deep FEF-deep LIP contact pairs), on state (wake vs anesthesia). For coherence estimates within or between layers of the same cortical layer, we included cortical area as a covariate. Thalamocortical comparisons were performed separately for each cortical area, and thus did not need this covariate. Similarly, cross-area corticocortical coherence, which was always computed between FEF and LIP, did not include this covariate. State was coded as a dichotomous variable (anesthesia = 1, wake = 2), and cortical area (Area) was coded as a centered, dichotomous variable (FEF = −0.5, LIP = 0.5) where applicable. Because spike-field coherence was calculated between individual neurons and derivatized LFPs, we included a random intercept by neuron (this inclusion changed neither the direction nor significance of the effects). Significant positive β_1_ parameters show frequency bands with increased power, coherence, or spike-field coherence during wakefulness relative to anesthesia. Significant negative β_1_ parameters show frequency bands with decreased power, coherence, or spike-field coherence during wakefulness relative to anesthesia.

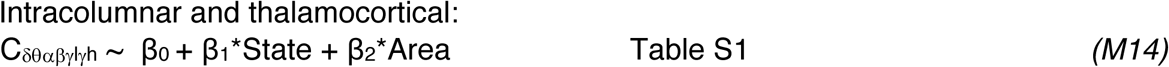

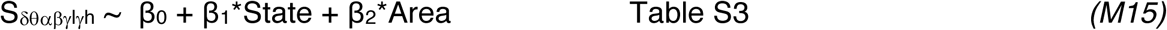

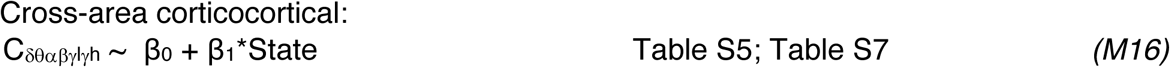

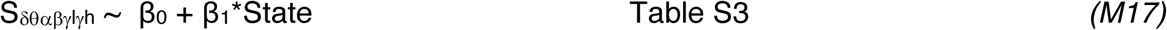

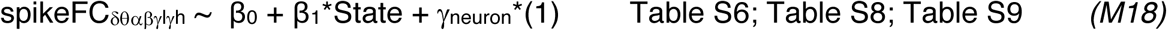

We limited all stimulation comparisons to anesthesia conditions without auditory stimuli where the stimulation frequency was 50 Hz and within the effective current range (120 - 200 μA). To test the effects of 50 Hz thalamic stimulation on power and coherence (Fig. 3, Fig. 4 and Fig. S5), we regressed change in power (S) and coherence (C) (stim – pre) averaged across different frequency bands (delta, alpha, theta, beta, low gamma, high gamma) and isolated to different electrode contact pairs of interest (in the case of coherence), on stimulation effectiveness (effective vs ineffective) including anesthetic and dose as covariates. For coherence estimates within or between layers of the same cortical layer, we included cortical area as a covariate. Cross-area corticocortical coherence, which was always computed between FEF and LIP, did not include this covariate. Anesthetic (Anes; isoflurane = −0.5, propofol = 0.5) and cortical area (Area; FEF = −0.5, LIP = 0.5), where applicable, were coded as centered, dichotomous variables. We coded dose as DoseCode. Significant positive β_1_ parameters show an interaction with stimulation epoch, where positive changes in power or coherence at the given frequency band are significantly larger for effective relative to ineffective stimulations. Significant negative β_1_ parameters show changes in power or coherence at the given frequency band that are significantly smaller for effective relative to ineffective stimulations. It was possible to get negative interactions even if power or coherence still increased during stimulation relative to the pre epoch.

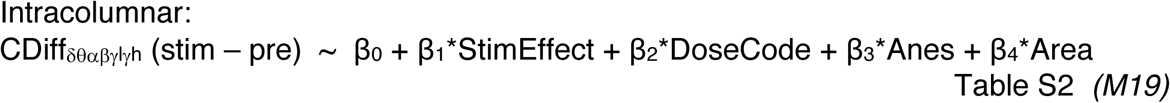

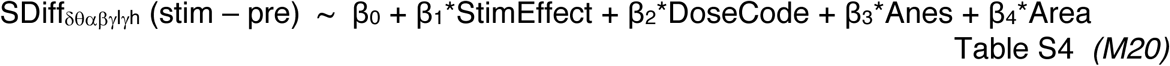

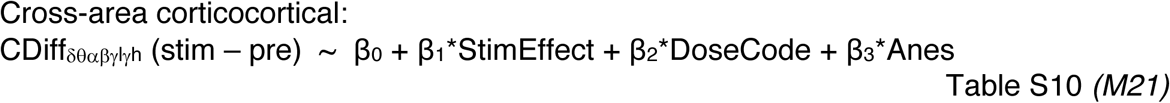

We considered effects consistent between (a) wake vs anesthesia and (b) effective vs ineffective stimulation comparisons (Fig. 3 and Fig. 4, gray shading) if both the StimEpoch effect from stimulation models (β_0_; stim – pre) were consistent in direction and significance to the beta parameter for the state effect in non-stimulation models (β_1_*State; wake – anesthesia). Such a finding indicates that the changes following stimulation-induced arousal are in the same direction as those found in the wake state over anesthesia. Additionally, the interaction term for stimulation data (β_1_*StimEffect) had to be significant, indicating that similarities were limited to the effective stimulation condition, and thus driven by arousal and not applied thalamic current in itself.

---

**Fig. S1.**
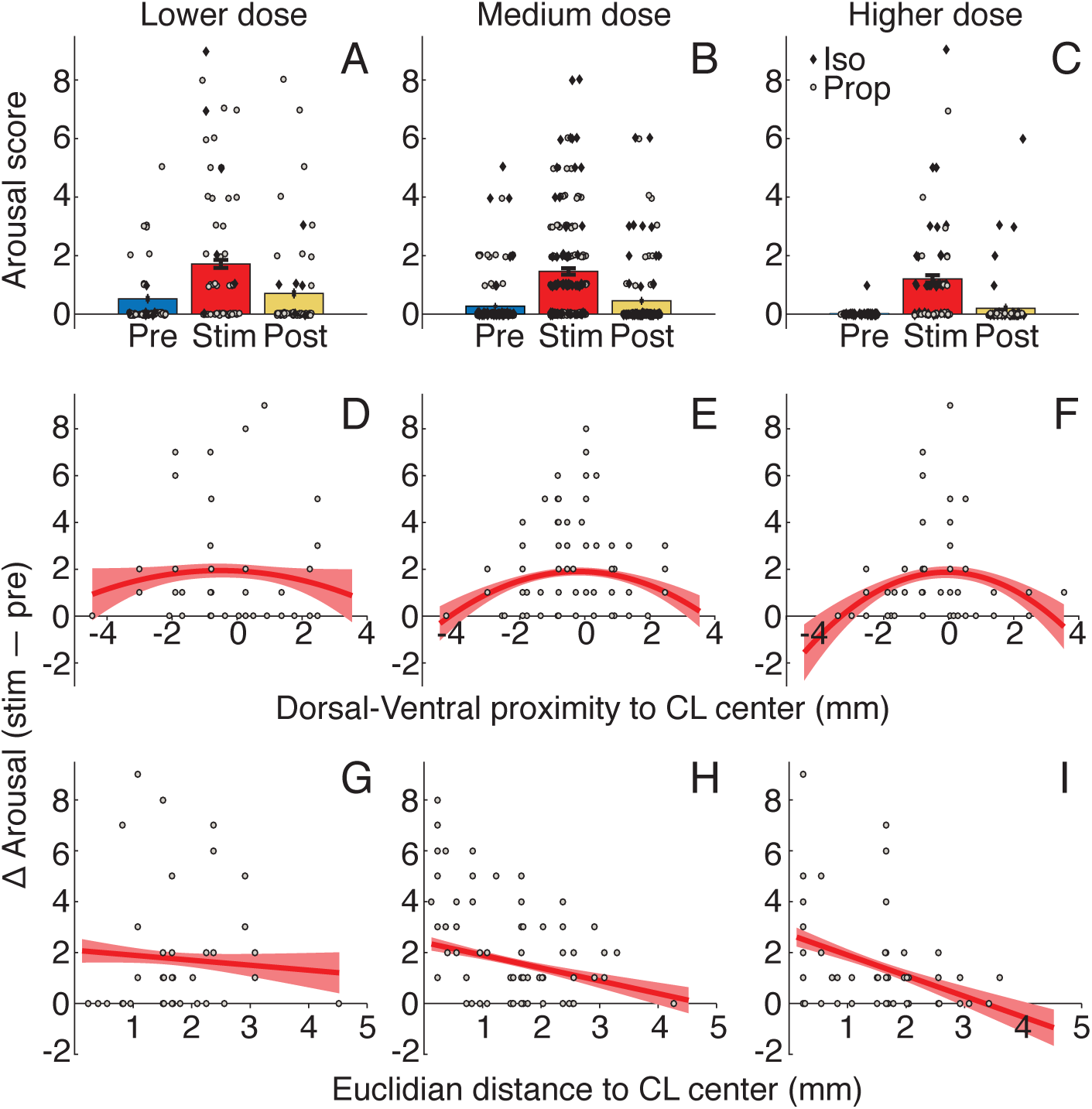
Stimulation effects on arousal did not significantly differ for different anesthetics or doses. (**A-C**) Population mean arousal score (±SE) from both monkeys prior to (blue), during (red), and after (yellow) thalamic stimulation at (**A**) lower, (**B**) medium and (**C**) higher anesthetic doses in our experimental range. Individual stimulation events under isoflurane (diamonds) and propofol (circles) shown. Stimulation effects on arousal occur irrespective of dose; although the interaction is not significant, effects are slightly stronger at lower doses in our experimental range. (**D-F**) Change in stimulation-induced arousal (stim – pre) as a function of dorsal-ventral distance from CL center at (**D**) lower, (**E**) medium and (**F**) higher doses in our experimental range. Circles represent individual stimulation events. Red curve indicates quadratic fit (±SE). Proximity to CL has a slightly stronger effect under higher doses of anesthesia (i.e., stimulating array may need to be closer to CL center to induce arousal at higher doses), but the interaction is not significant. (**G-I**) Change in stimulation-induced arousal (stim – pre) as a function of Euclidian distance from CL center at (**G**) lower, (**H**) medium and (**I**) higher doses in our experimental range. Circles represent individual stimulation events. Red curve indicates linear fit (±SE). Proximity to CL is significantly predictive of arousal score regardless of dose. The effect is slightly stronger under higher doses of anesthesia, but the interaction is not significant.

**Fig. S2.**
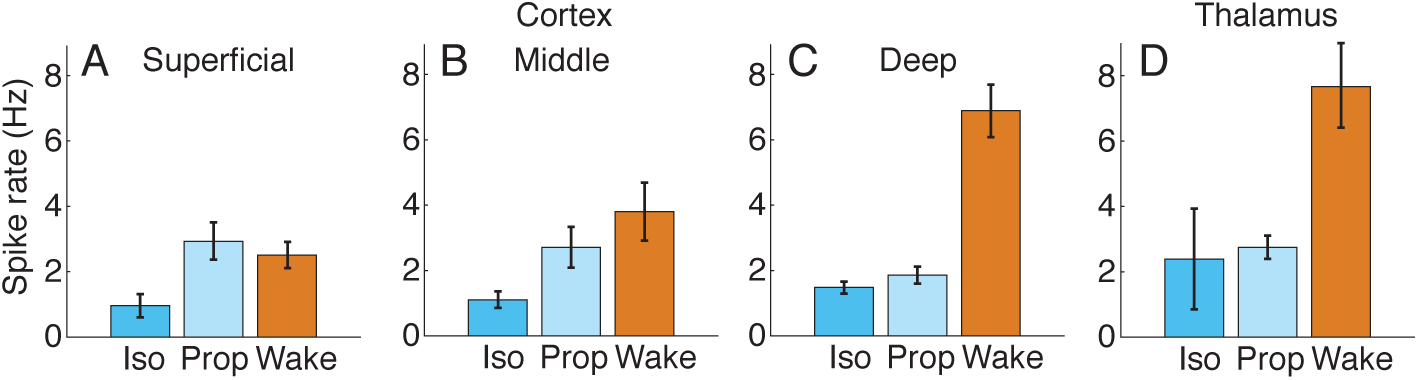
Propofol and isoflurane had similar effects on spiking activity. (**A-D**) Spike rate of neurons recorded during isoflurane (Iso, blue), propofol (Prop, light blue) and wakefulness (Wake, orange) for (**A**) superficial cortical, (**B**) middle cortical, (**C**) deep cortical and (**D**) thalamic neurons. No significant differences were found between anesthetics.

**Fig. S3.**
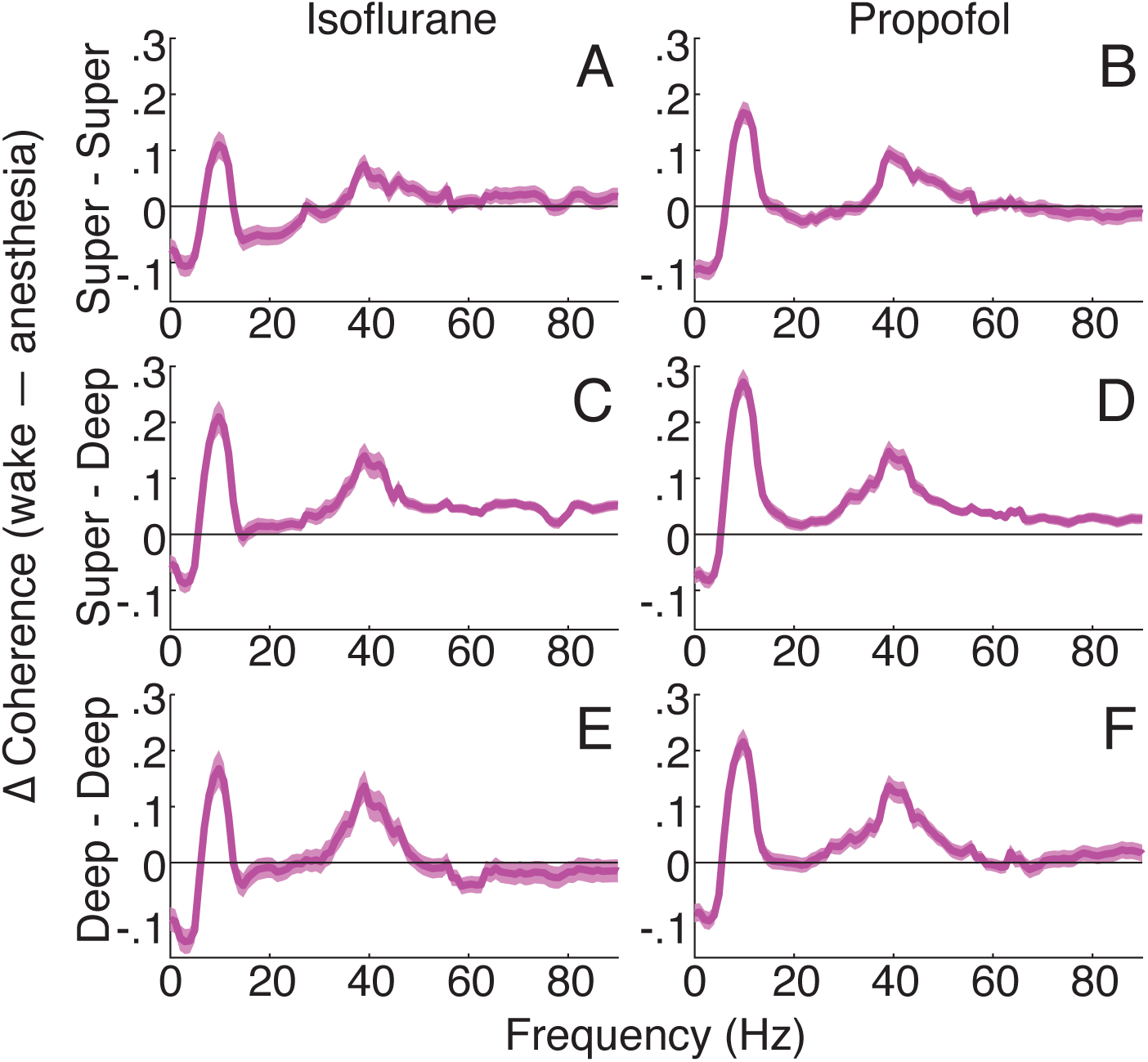
Propofol and isoflurane had similar influences on intracolumnar interactions. (**A-F**) Population coherence difference between wakefulness and anesthesia. Positive when wake > anesthesia. Error bars indicate ±SE of T-tests at each frequency. Average of all contact pairs (cortical areas combined) for: superficial cortical layers under (**A**) isoflurane and (**B**) propofol; between superficial and deep layers under (**C**) isoflurane and (**D**) propofol; and for deep layers under (**E**) isoflurane and (**F**) propofol.

**Fig. S4.**
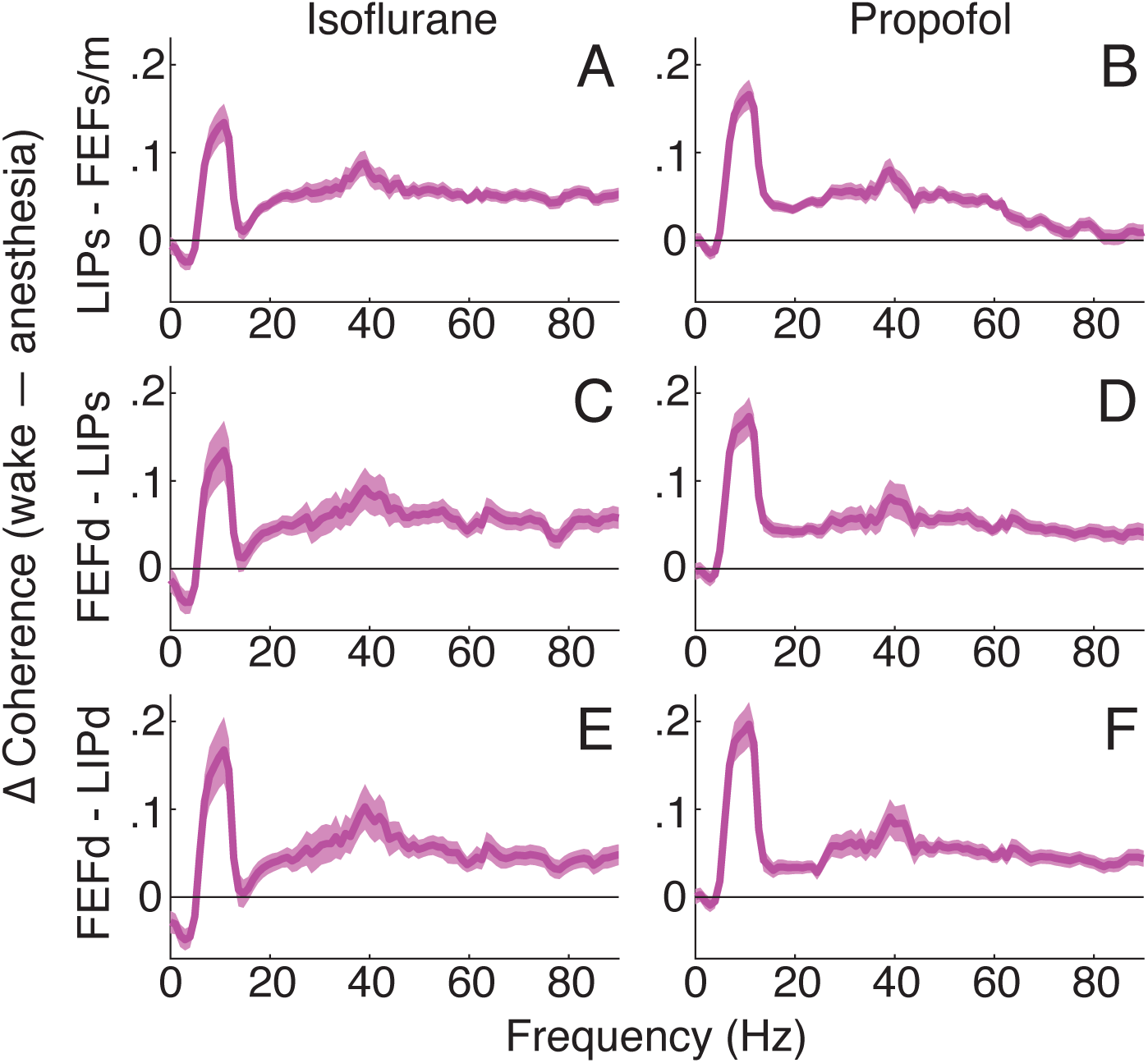
Propofol and isoflurane had similar influences on cross-area interactions. (**A-F**) Population average coherence difference between wakefulness and anesthesia. Positive when wake > anesthesia. Error bars indicate ±SE of T-tests at each frequency. Average of all contact pairs for: superficial LIP and superficial/mid FEF under (**A**) isoflurane and (**B**) propofol; deep FEF and superficial LIP under (**C**) isoflurane and (**D**) propofol; and deep FEF and deep LIP under (**E**) isoflurane and (**F**) propofol.

**Fig. S5.**
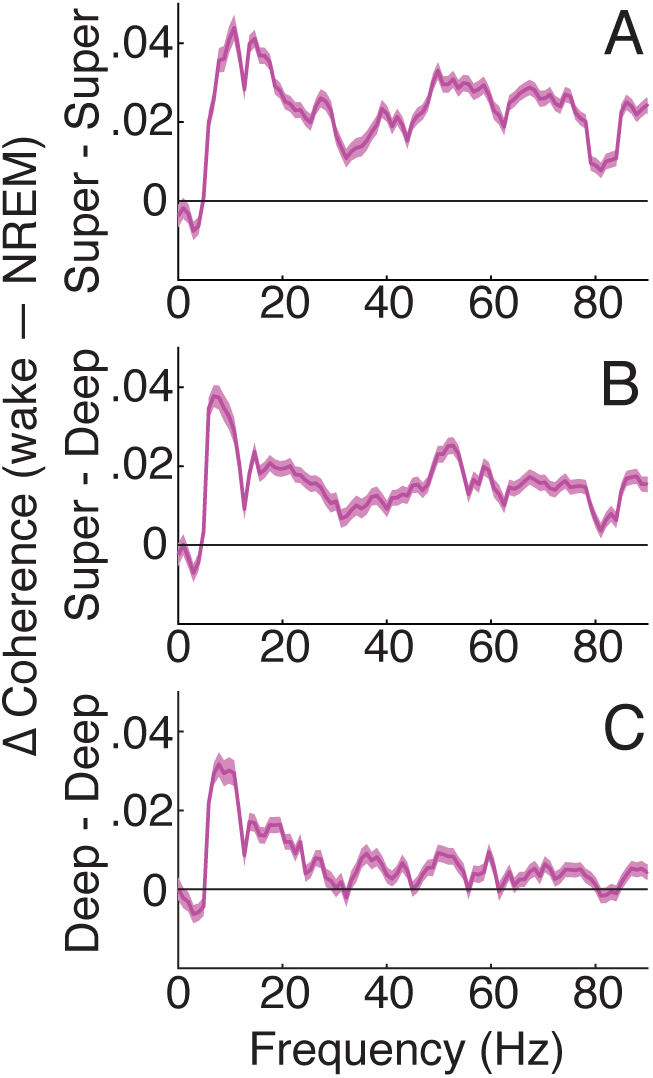
Light NREM sleep had qualitatively similar influences on intracolumnar interactions as anesthesia. (**A-C**) Population average coherence difference between wakefulness and sleep. Positive when wake > sleep. Error bars indicate ±SE. Average of all contact pairs (cortical areas combined) for: (**A**) superficial cortical layers; (**B**) between superficial and deep layers; and (**C**) deep layers.

**Fig. S6.**
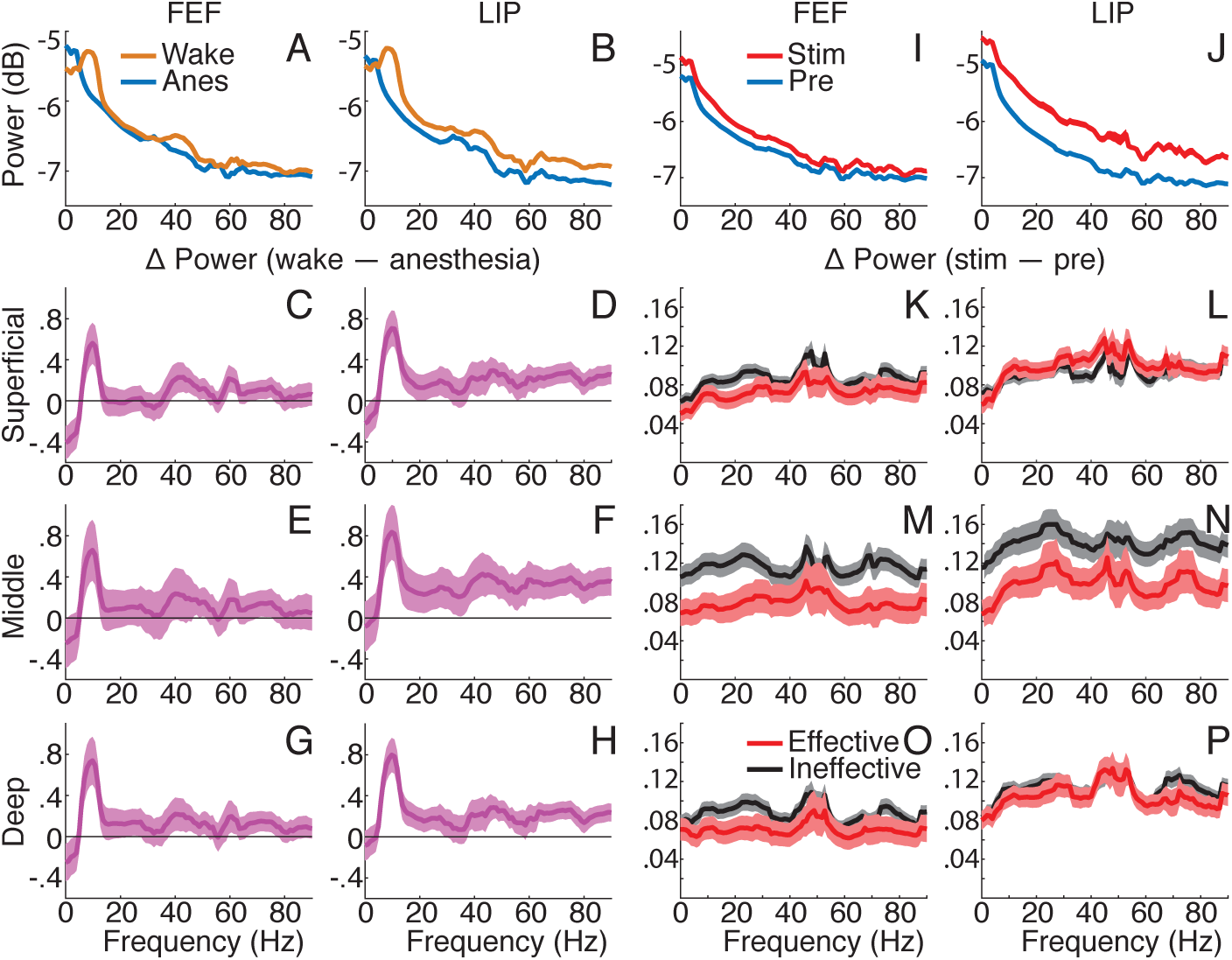
Cortical power correlates poorly with arousal. Effects were not consistent between state (wake vs anesthesia) and stimulation (effective vs ineffective), and thus power is unlikely to be key contributor to the NCC. (**A**) Population FEF and (**B**) LIP power spectra for wakefulness (orange) and anesthesia (blue). Average of all contacts across cortical layers. Line thickness indicates ±SE. (**C-H**) Population power difference between wakefulness and anesthesia. Positive when wake > anesthesia. Error bars indicate ±SE of T-tests at each frequency. Average of all contacts for: superficial (**C**) FEF and (**D**) LIP; middle (**E**) FEF and (**F**) LIP; deep (**G**) FEF and (**H**) LIP. (**I**) Population FEF and (**J**) LIP power under anesthesia prior to (blue) and during effective stimulation (red). Average across all cortical layers. Line thickness indicates ±SE. (**K-P**) Population power difference between effective (red) and ineffective (black) stimulations. Positive when stim > pre. Error bars indicate ±SE. Average of all contacts for: superficial (**K**) FEF and (**L**) LIP; middle (**M**) FEF and (**N**) LIP; deep (**O**) FEF and (**P**) LIP.

**Fig. S7.**
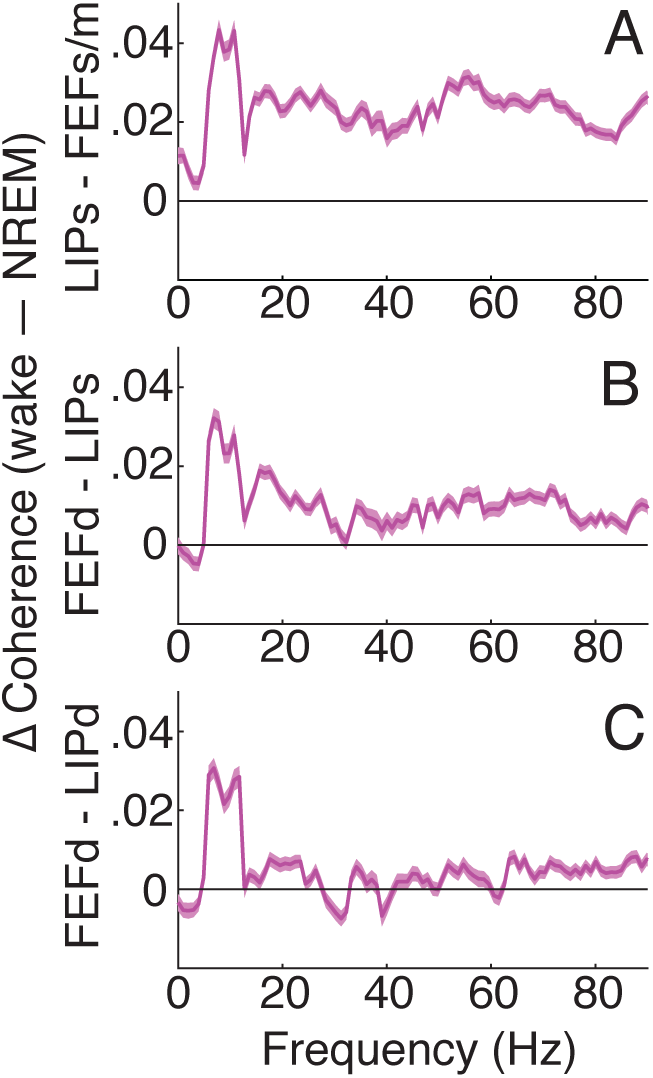
Light NREM sleep had qualitatively similar influences on cross-area interactions as anesthesia. (**A-C**) Population average coherence difference between wakefulness and sleep. Positive when wake > sleep. Error bars indicate ±SE. Average of all contact pairs for: (**A**) superficial LIP and superficial/mid FEF; (**B**) deep FEF and superficial LIP; and (**C**) deep FEF and deep LIP.

**Fig. S8.**
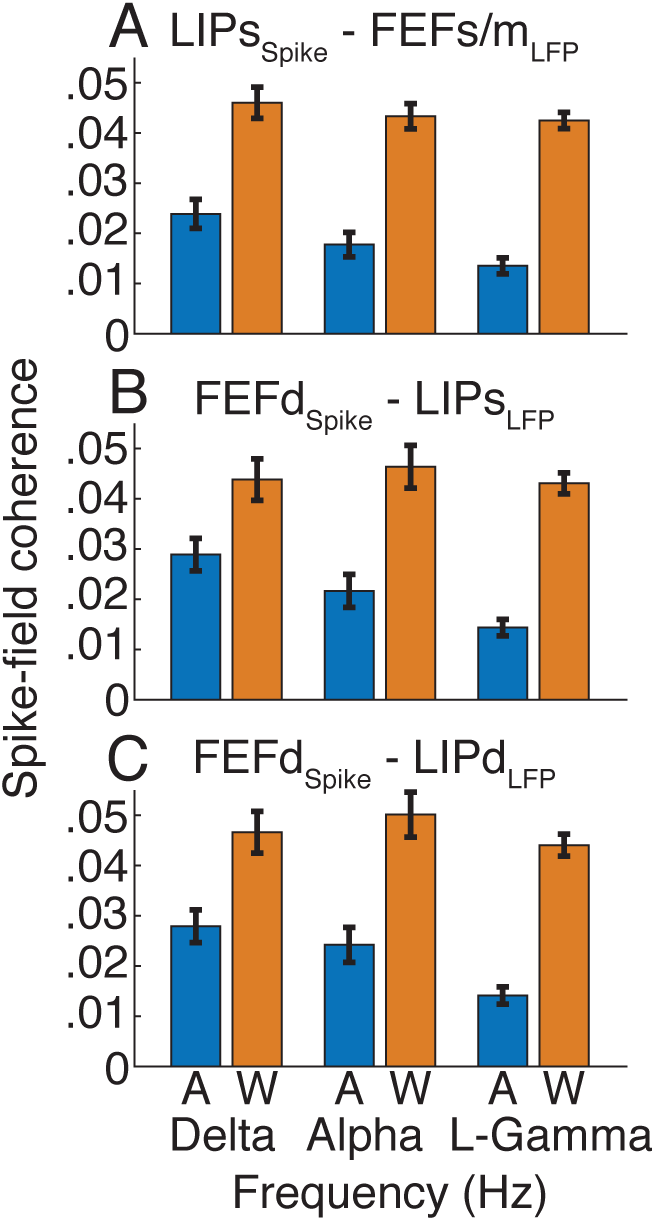
Spike-field coherence of corticocortical pathways reduced during anesthesia. (**A-C**) Spike-field coherence (±SE) at delta (0-4 Hz), alpha (8-15 Hz) and low gamma (30-60 Hz) frequencies during wakefulness (W, orange) and anesthesia (A, blue). Average of all contact pairs for: (**A**) superficial LIP spikes and superficial/middle FEF LFPs; (**B**) deep FEF spikes and superficial LIP LFPs; and (**C**) deep FEF spikes and deep LIP LFPs.

**Fig. S9.**
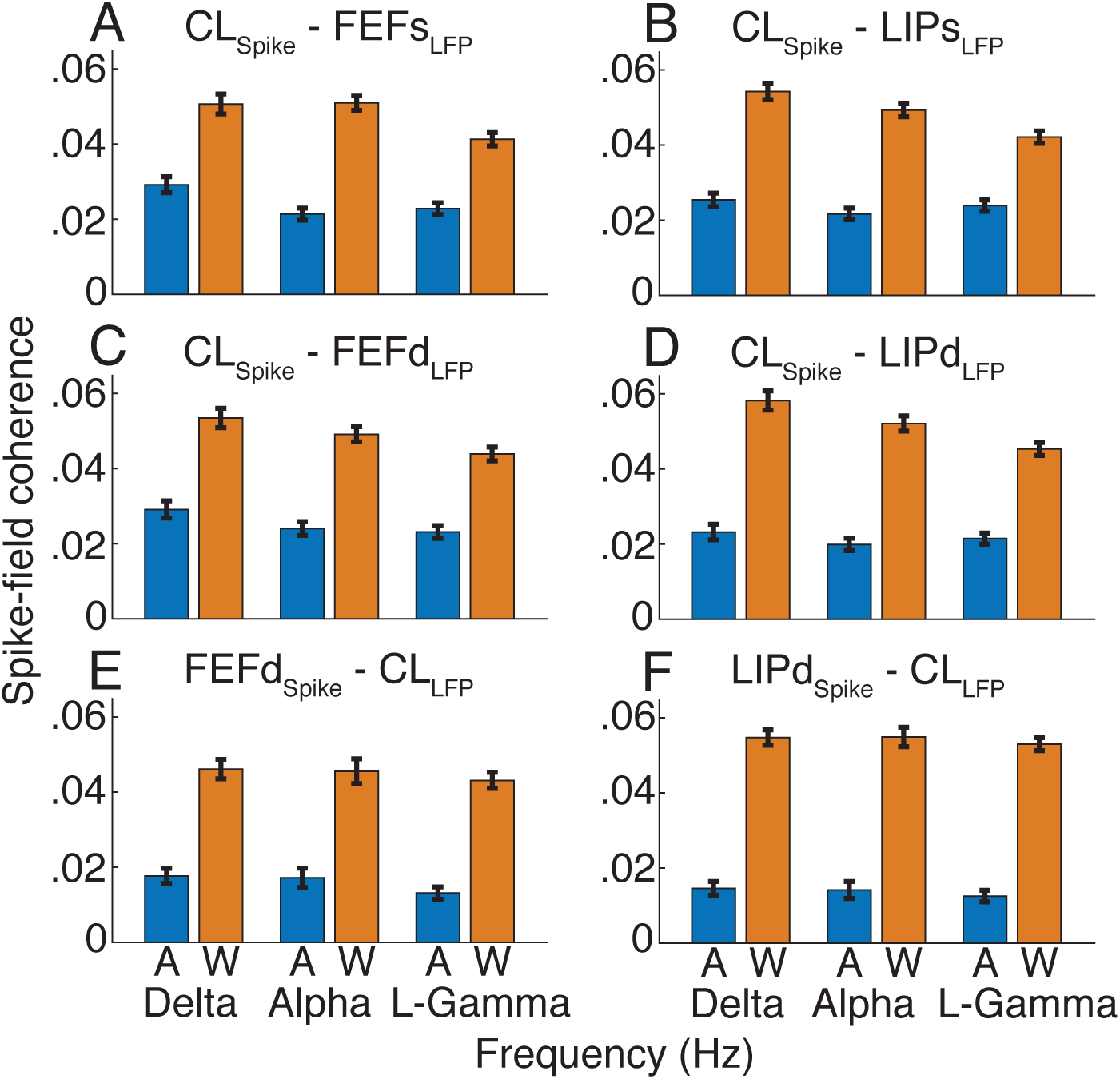
Spike-field coherence of thalamocortical pathways reduced during anesthesia. (**A-F**) Spike-field coherence (±SE) at delta (0-4 Hz), alpha (8-15 Hz) and low gamma (30-60 Hz) during wake (W, orange) and anesthesia (A, blue). Average of all contact pairs for: CL spikes to superficial LFPs in (**A**) FEF and (**B**) LIP; CL spikes to deep LFPs in (**C**) FEF and (**D**) LIP; and deep cortical spikes in (**E**) FEF and (**F**) LIP to thalamic LFPs.

**Table S1.** Statistical results for intracolumnar cortical coherence in wakefulness and anesthesia. Analyses with model 14 performed for: delta (δ) = 0-4 Hz; theta (θ) = 4-8 Hz; alpha (α) = 8-15 Hz; beta (β) = 15-30; low gamma (γ_l_) = 30-60 Hz; and high gamma (γ_h_) = 60-90 Hz for coherence pairs within/between different layers of cortical areas. Reported statistics are the slope (β_1_), T statistic (T) and Holm’s adjusted p-value (p_adj_) for the parameter of interest (β_1_*State). Significant effects (p < 0.05) show frequency bands where coherence is significantly different for wakefulness relative to anesthesia (Anes).

| M14 | Data | Band | $\beta_1$ | T | $p_{adj}$ |
| --- | --- | --- | --- | --- | --- |
| $C \sim \beta_0 + \beta_1 * \text{State} + \beta_2 * \text{Area}$ | **Wake v Anes**<br>**Intracolumnar**<br>**Sup ↔ Sup**<br>N<sub>Wake</sub> = 4326<br>N<sub>Anes</sub> = 5959<br>DF = 10282 | $\delta$ | -0.104 | -28.08 | $<1.0 \times 10^{-10}$ |
| | | $\theta$ | 0.002 | 0.43 | 1.000 |
| | | $\alpha$ | 0.076 | 20.86 | $<1.0 \times 10^{-10}$ |
| | | $\beta$ | -0.028 | -10.23 | $<1.0 \times 10^{-10}$ |
| | | $\gamma_l$ | 0.014 | 5.91 | $1.7 \times 10^{-8}$ |
| | | $\gamma_h$ | -0.003 | -1.07 | 0.937 |
| | **Wake v Anes**<br>**Intracolumnar**<br>**Sup ↔ Deep**<br>N<sub>Wake</sub> = 4196<br>N<sub>Anes</sub> = 4529<br>DF = 8722 | $\delta$ | -0.073 | -22.78 | $<1.0 \times 10^{-10}$ |
| | | $\theta$ | 0.086 | 23.29 | $<1.0 \times 10^{-10}$ |
| | | $\alpha$ | 0.154 | 41.79 | $<1.0 \times 10^{-10}$ |
| | | $\beta$ | 0.020 | 10.05 | $<1.0 \times 10^{-10}$ |
| | | $\gamma_l$ | 0.054 | 25.12 | $<1.0 \times 10^{-10}$ |
| | | $\gamma_h$ | 0.034 | 23.02 | $<1.0 \times 10^{-10}$ |
| | **Wake v Anes**<br>**Intracolumnar**<br>**Deep ↔ Deep**<br>N<sub>Wake</sub> = 4378<br>N<sub>Anes</sub> = 4316<br>DF = 8691 | $\delta$ | -0.102 | -26.66 | $<1.0 \times 10^{-10}$ |
| | | $\theta$ | 0.053 | 12.70 | $<1.0 \times 10^{-10}$ |
| | | $\alpha$ | 0.105 | 26.04 | $<1.0 \times 10^{-10}$ |
| | | $\beta$ | -0.002 | -0.62 | 1.000 |
| | | $\gamma_l$ | 0.036 | 11.87 | $<1.0 \times 10^{-10}$ |
| | | $\gamma_h$ | -0.004 | -1.19 | 0.937 |

**Table S2.** Statistical results for intracolumnar cortical coherence during effective and ineffective stimulations. Analyses with model 19 performed for: delta (δ) = 0-4 Hz; theta (θ) = 4-8 Hz; alpha (α) = 8-15 Hz; beta(β) = 15-30 Hz, low Gamma(γ_l_) = 30-60 Hz; and high gamma(γ_h_) = 60-90 Hz for coherence pairs within/between different layers of cortical areas. Reported statistics are the slope (β_1_), T statistic (T) and Holm’s adjusted p-value (p_adj_) for the parameter of interest (β_1_*StimEffect). Significant interactions (p < 0.05) show frequency bands where changes induced by stimulation are significantly different for effective relative to ineffective stimulations.

| M19 | Data | Band | $\beta_1$ | T | $p_{adj}$ |
| --- | --- | --- | --- | --- | --- |
| CDiff (stim – pre) ~ $\beta_0 + \beta_1 * \text{StimEffect} + \beta_2 * \text{DoseCode} + \beta_3 * \text{Anes} + \beta_4 * \text{Area}$ | **Stimulation Intracolumnar Sup ⇔ Sup**<br>N<sub>Effect</sub> = 845<br>N<sub>Ineffect</sub> = 1544<br>DF = 2384 | $\delta$ | -0.002 | -0.30 | 1.000 |
| | | $\theta$ | 0.009 | 1.18 | 1.000 |
| | | $\alpha$ | 0.017 | 2.44 | 0.119 |
| | | $\beta$ | -0.004 | -0.85 | 1.000 |
| | | $\gamma_l$ | 0.020 | 5.24 | $1.8 \times 10^{-6}$ |
| | | $\gamma_h$ | 0.003 | 0.77 | 1.000 |
| | **Stimulation Intracolumnar Sup ⇔ Deep**<br>N<sub>Effect</sub> = 826<br>N<sub>Ineffect</sub> = 1805<br>DF = 2626 | $\delta$ | 0.006 | 0.82 | 1.000 |
| | | $\theta$ | 0.032 | 4.83 | $1.3 \times 10^{-6}$ |
| | | $\alpha$ | 0.045 | 7.31 | $<1.0 \times 10^{-10}$ |
| | | $\beta$ | 0.021 | 5.42 | $7.3 \times 10^{-6}$ |
| | | $\gamma_l$ | 0.032 | 10.45 | $<1.0 \times 10^{-10}$ |
| | | $\gamma_h$ | 0.023 | 7.50 | $<1.0 \times 10^{-10}$ |
| | **Stimulation Intracolumnar Deep ⇔ Deep**<br>N<sub>Effect</sub> = 699<br>N<sub>Ineffect</sub> = 1484<br>DF = 2178 | $\delta$ | 0.069 | 7.75 | $<1.0 \times 10^{-10}$ |
| | | $\theta$ | 0.070 | 9.04 | $<1.0 \times 10^{-10}$ |
| | | $\alpha$ | 0.083 | 11.79 | $<1.0 \times 10^{-10}$ |
| | | $\beta$ | 0.034 | 7.93 | $<1.0 \times 10^{-10}$ |
| | | $\gamma_l$ | 0.008 | 2.21 | 0.190 |
| | | $\gamma_h$ | 0.005 | 1.58 | 0.681 |

**Table S3.**
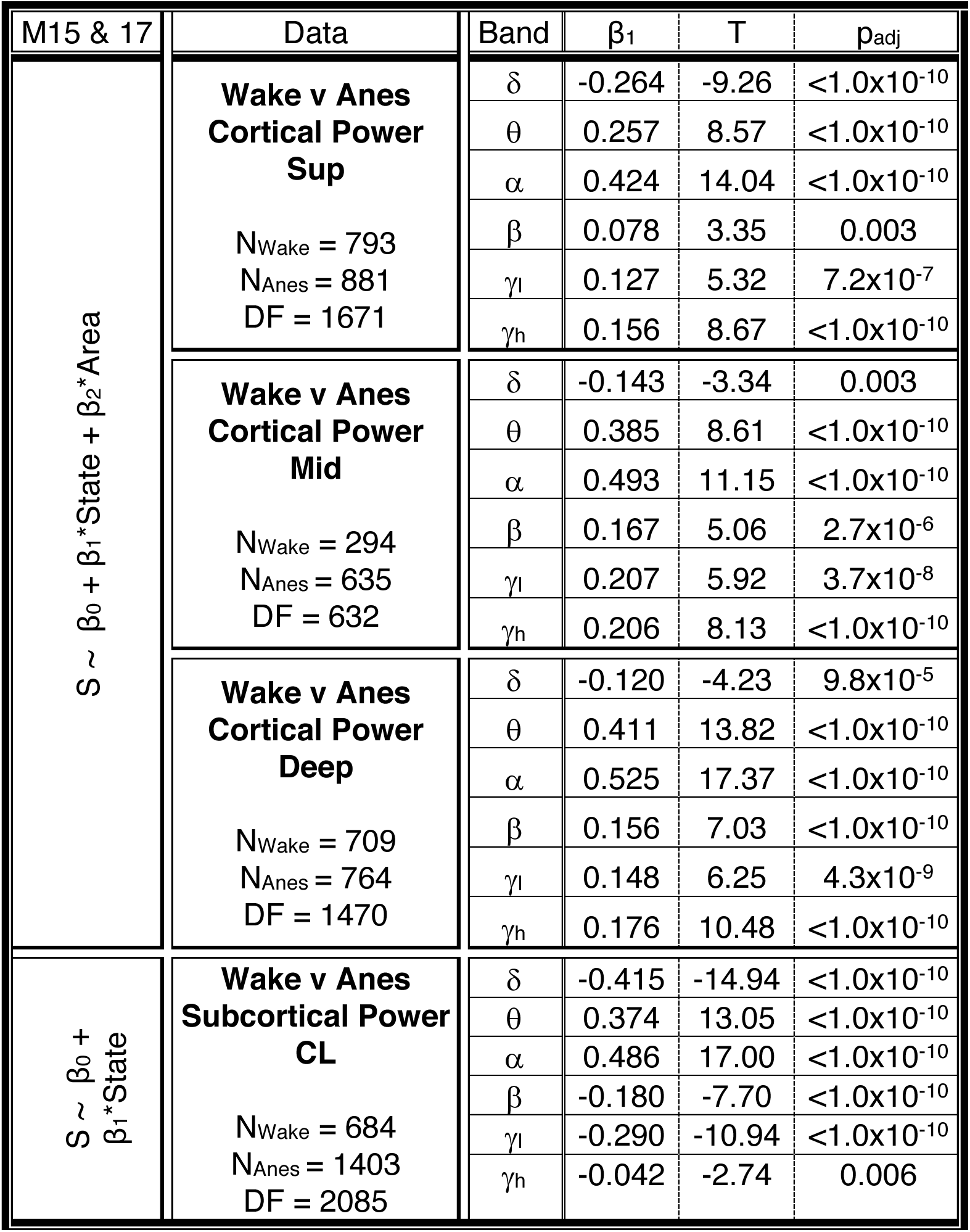
Statistical results for cortical and thalamic power in wakefulness and anesthesia. Analyses with model 15 and 17 performed for: delta (δ) = 0-4 Hz; theta (θ) = 4-8 Hz; alpha (α) = 8-15 Hz; beta (β) = 15-30 Hz; low gamma (γ_l_) = 30-60 Hz; and high gamma (γ_h_) = 60-90 Hz for different layers of cortical areas and for thalamus. Reported statistics are the slope (β_1_), T statistic (T) and Holm’s adjusted p-value (p_adj_) for the parameter of interest (β_1_*State). Significant effects (p < 0.05) show frequency bands where power is significantly different for wakefulness relative to anesthesia (Anes).

**Table S4.**
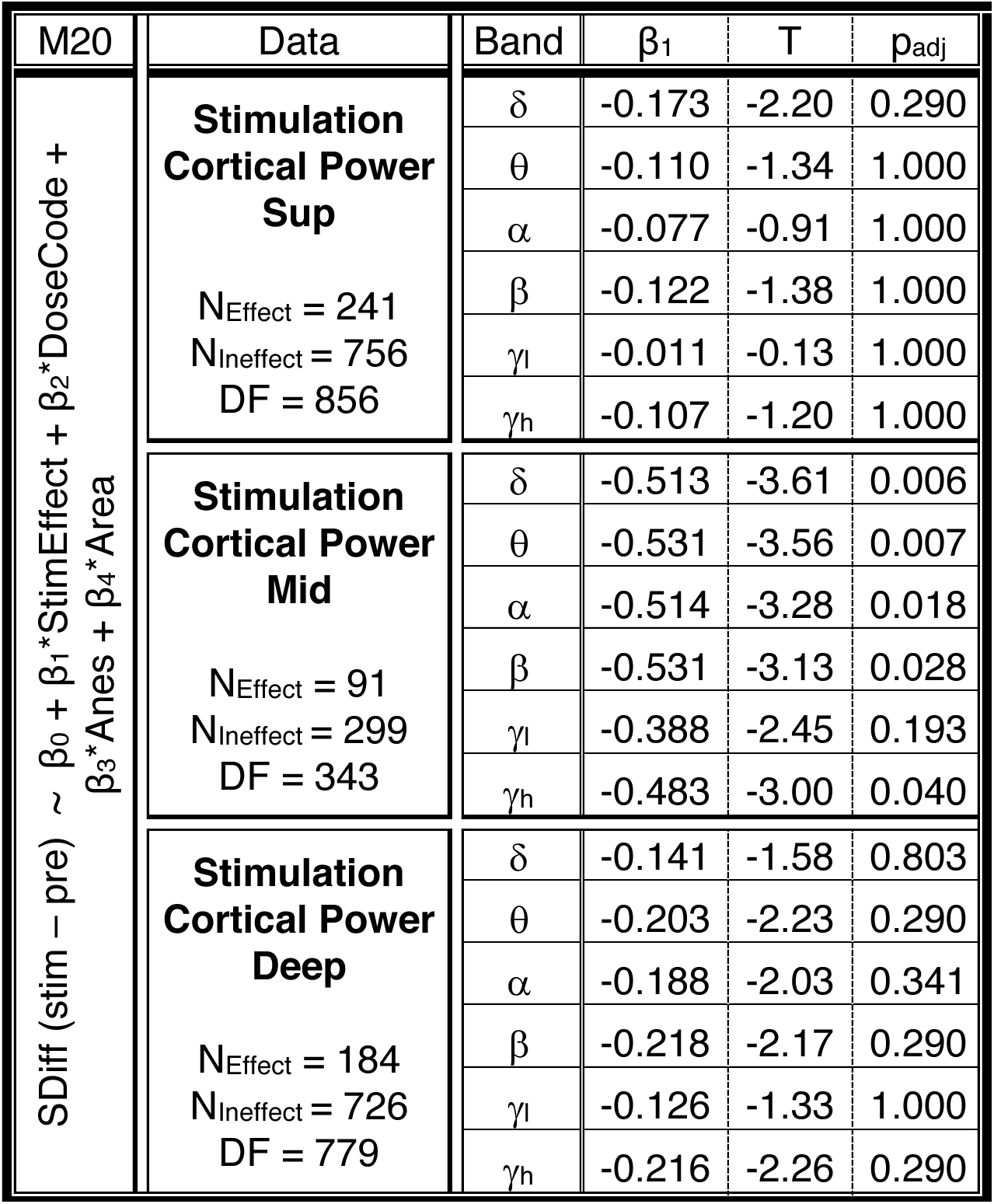
Statistical results for cortical power during effective and ineffective stimulations. Analyses with model 20 performed for: delta (δ) = 0-4 Hz; theta (θ) = 4-8 Hz; alpha (α) = 8-15 Hz; beta (β) = 15-30 Hz; low gamma (γ_l_) = 30-60 Hz; and high gamma (γ_h_) = 60-90 Hz for contacts within different cortical layers. Reported statistics are the slope (β_1_), T statistic (T) and Holm’s adjusted p-value (p_adj_) for the parameter of interest (β_1_*StimEffect). Significant interactions (p < 0.05) show frequency bands where changes induced by stimulation are significantly different for effective relative to ineffective stimulations.

| M20 | Data | Band | $\beta_1$ | T | $p_{adj}$ |
| --- | --- | --- | --- | --- | --- |
| $S_{Diff} (stim - pre) \sim \beta_0 + \beta_1 * StimEffect + \beta_2 * DoseCode + \beta_3 * Anes + \beta_4 * Area$ | **Stimulation Cortical Power Sup**<br>$N_{Effect} = 241$<br>$N_{Ineffect} = 756$<br>$DF = 856$ | $\delta$ | -0.173 | -2.20 | 0.290 |
| | | $\theta$ | -0.110 | -1.34 | 1.000 |
| | | $\alpha$ | -0.077 | -0.91 | 1.000 |
| | | $\beta$ | -0.122 | -1.38 | 1.000 |
| | | $\gamma_l$ | -0.011 | -0.13 | 1.000 |
| | | $\gamma_h$ | -0.107 | -1.20 | 1.000 |
| | **Stimulation Cortical Power Mid**<br>$N_{Effect} = 91$<br>$N_{Ineffect} = 299$<br>$DF = 343$ | $\delta$ | -0.513 | -3.61 | 0.006 |
| | | $\theta$ | -0.531 | -3.56 | 0.007 |
| | | $\alpha$ | -0.514 | -3.28 | 0.018 |
| | | $\beta$ | -0.531 | -3.13 | 0.028 |
| | | $\gamma_l$ | -0.388 | -2.45 | 0.193 |
| | | $\gamma_h$ | -0.483 | -3.00 | 0.040 |
| | **Stimulation Cortical Power Deep**<br>$N_{Effect} = 184$<br>$N_{Ineffect} = 726$<br>$DF = 779$ | $\delta$ | -0.141 | -1.58 | 0.803 |
| | | $\theta$ | -0.203 | -2.23 | 0.290 |
| | | $\alpha$ | -0.188 | -2.03 | 0.341 |
| | | $\beta$ | -0.218 | -2.17 | 0.290 |
| | | $\gamma_l$ | -0.126 | -1.33 | 1.000 |
| | | $\gamma_h$ | -0.216 | -2.26 | 0.290 |

**Table S5.** Statistical results for cross-area corticocortical LFP-LFP coherence in wakefulness and anesthesia. Analyses with model 16 performed for: delta (δ) = 0-4 Hz; theta (θ) = 4- 8 Hz; alpha (α) = 8-15 Hz; beta (β) = 15-30 Hz; low gamma (γ_l_) = 30-60 Hz; and high gamma (γ_h_) = 60-90 Hz for different pairs of contacts between FEF and LIP (deep (D), superficial (S), and middle (M) layers). Reported statistics are the slope (β_1_), T statistic (T) and Holm’s adjusted p-value (p_adj_) for the parameter of interest (State). Significant effects (p < 0.05) show frequency bands where coherence is significantly different for wakefulness relative to anesthesia.

| M16 | Data | Band | $\beta_1$ | T | $p_{adj}$ |
| --- | --- | --- | --- | --- | --- |
| $C \sim \beta_0 + \beta_1 * \text{State}$ | **Wake v Anes**<br>**Cross-area corticocortical**<br>**LIP<sub>s</sub> ↔ FEF<sub>S/M</sub>**<br>N<sub>Wake</sub> = 3934<br>N<sub>Anes</sub> = 3664<br>DF = 7524 | $\delta$ | -0.012 | -6.91 | $<1.0 \times 10^{-10}$ |
| | | $\theta$ | 0.070 | 32.30 | $<1.0 \times 10^{-10}$ |
| | | $\alpha$ | 0.103 | 41.80 | $<1.0 \times 10^{-10}$ |
| | | $\beta$ | 0.045 | 50.98 | $<1.0 \times 10^{-10}$ |
| | | $\gamma_l$ | 0.061 | 46.64 | $<1.0 \times 10^{-10}$ |
| | | $\gamma_h$ | 0.027 | 19.48 | $<1.0 \times 10^{-10}$ |
| | **Wake v Anes**<br>**Cross-area corticocortical**<br>**FEF<sub>D</sub> ↔ LIP<sub>s</sub>**<br>N<sub>Wake</sub> = 2634<br>N<sub>Anes</sub> = 1816<br>DF = 4410 | $\delta$ | -0.015 | -7.64 | $<1.0 \times 10^{-10}$ |
| | | $\theta$ | 0.092 | 28.34 | $<1.0 \times 10^{-10}$ |
| | | $\alpha$ | 0.118 | 31.12 | $<1.0 \times 10^{-10}$ |
| | | $\beta$ | 0.039 | 30.06 | $<1.0 \times 10^{-10}$ |
| | | $\gamma_l$ | 0.071 | 41.86 | $<1.0 \times 10^{-10}$ |
| | | $\gamma_h$ | 0.044 | 32.50 | $<1.0 \times 10^{-10}$ |
| | **Wake v Anes**<br>**Cross-area corticocortical**<br>**FEF<sub>D</sub> ↔ LIP<sub>D</sub>**<br>N<sub>Wake</sub> = 2083<br>N<sub>Anes</sub> = 1947<br>DF = 3968 | $\delta$ | -0.013 | -6.04 | $1.6 \times 10^{-9}$ |
| | | $\theta$ | 0.082 | 27.24 | $<1.0 \times 10^{-10}$ |
| | | $\alpha$ | 0.108 | 31.65 | $<1.0 \times 10^{-10}$ |
| | | $\beta$ | 0.047 | 39.02 | $<1.0 \times 10^{-10}$ |
| | | $\gamma_l$ | 0.069 | 41.25 | $<1.0 \times 10^{-10}$ |
| | | $\gamma_h$ | 0.047 | 33.31 | $<1.0 \times 10^{-10}$ |

**Table S6.** Statistical results for cross-area corticocortical spike-field coherence in wakefulness and anesthesia. Analyses with model 18 performed for: delta (δ) = 0-4 Hz; theta (θ) = 4-8 Hz; alpha (α) = 8-15 Hz; beta (β) = 15-30 Hz; low gamma (γ_l_) = 30-60 Hz; and high gamma (γ_h_) = 60-90 Hz for different contact pairs between FEF and LIP in different layers. Spike-field coherence between superficial LIP spikes and superficial/mid FEF LFPs (top), deep FEF spikes and superficial LIP LFPs (middle), and deep FEF spikes and deep LIP LFPs (bottom). Reported statistics are the slope (β_1_), F statistic (F) and Holm’s adjusted p-value (p_adj_) for the parameter of interest (β_1_*State). Significant effects (p < 0.05) show bands where coherence is significantly different for wakefulness relative to anesthesia.

| M18 | Data | Band | $\beta_1$ | F | $p_{adj}$ |
| --- | --- | --- | --- | --- | --- |
| spikeFC $\sim \beta_0 + \beta_1 * \text{State} + \gamma_{\text{neuron}}^*(1)$ | **Wake v Anes**<br>**Cross-area**<br>**corticocortical**<br>**LIP<sub>s</sub> ↔ FEF<sub>S/M</sub>**<br>N<sub>Wake</sub> = 30<br>N<sub>Anes</sub> = 30<br>DF = 57.43 | $\delta$ | 0.022 | 27.26 | $1.8 \times 10^{-5}$ |
| | | $\theta$ | 0.024 | 42.17 | $1.7 \times 10^{-7}$ |
| | | $\alpha$ | 0.026 | 53.29 | $8.4 \times 10^{-9}$ |
| | | $\beta$ | 0.027 | 154.39 | $<1.0 \times 10^{-10}$ |
| | | $\gamma_l$ | 0.029 | 161.93 | $<1.0 \times 10^{-10}$ |
| | | $\gamma_h$ | 0.030 | 155.71 | $<1.0 \times 10^{-10}$ |
| | **Wake v Anes**<br>**Cross-area**<br>**corticocortical**<br>**FEF<sub>D</sub> ↔ LIP<sub>s</sub>**<br>N<sub>Wake</sub> = 44<br>N<sub>Anes</sub> = 74<br>DF = 114.08 | $\delta$ | 0.015 | 8.04 | 0.005 |
| | | $\theta$ | 0.026 | 19.76 | $6.4 \times 10^{-5}$ |
| | | $\alpha$ | 0.025 | 21.23 | $6.3 \times 10^{-5}$ |
| | | $\beta$ | 0.027 | 71.84 | $<1.0 \times 10^{-10}$ |
| | | $\gamma_l$ | 0.029 | 116.59 | $<1.0 \times 10^{-10}$ |
| | | $\gamma_h$ | 0.028 | 123.53 | $<1.0 \times 10^{-10}$ |
| | **Wake v Anes**<br>**Cross-area**<br>**corticocortical**<br>**FEF<sub>D</sub> ↔ LIP<sub>D</sub>**<br>N<sub>Wake</sub> = 45<br>N<sub>Anes</sub> = 75<br>DF = 115.19 | $\delta$ | 0.019 | 12.59 | 0.001 |
| | | $\theta$ | 0.031 | 20.33 | $6.4 \times 10^{-5}$ |
| | | $\alpha$ | 0.026 | 20.78 | $6.4 \times 10^{-5}$ |
| | | $\beta$ | 0.031 | 66.96 | $<1.0 \times 10^{-10}$ |
| | | $\gamma_l$ | 0.030 | 114.38 | $<1.0 \times 10^{-10}$ |
| | | $\gamma_h$ | 0.029 | 145.27 | $<1.0 \times 10^{-10}$ |

**Table S7.** Statistical results for cross-area thalamocortical LFP-LFP coherence. Analyses with model 16 performed for: delta (δ) = 0-4 Hz; theta (θ) = 4-8 Hz; alpha (α) = 8-15 Hz; beta (β) = 15-30 Hz; low gamma (γ_l_) = 30-60 Hz; and high gamma (γ_h_) = 60-90 Hz for different contact pairs between thalamus and superficial (S) or deep (D) cortical layers in FEF or LIP. Reported statistics are the slope (β_1_), T statistic (T) and Holm’s adjusted p-value (p_adj_) for the parameter of interest (β_1_*State). Significant effects (p < 0.05) show frequency bands where coherence is significantly different for wakefulness relative to anesthesia.

| M16 | Data | Band | $\beta_1$ | T | $p_{adj}$ |
| --- | --- | --- | --- | --- | --- |
| $C \sim \beta_0 + \beta_1 * \text{State}$ | **Wake v Anes**<br>**Cross-area**<br>**CL ⇔ FEF<sub>s</sub>**<br>N<sub>Wake</sub> = 3446<br>N<sub>Anes</sub> = 9315<br>DF = 12531 | $\delta$ | 0.030 | 20.68 | <1.0x10<sup>-10</sup> |
| | | $\theta$ | 0.084 | 59.83 | <1.0x10<sup>-10</sup> |
| | | $\alpha$ | 0.097 | 64.15 | <1.0x10<sup>-10</sup> |
| | | $\beta$ | 0.045 | 57.72 | <1.0x10<sup>-10</sup> |
| | | $\gamma_l$ | 0.065 | 66.28 | <1.0x10<sup>-10</sup> |
| | | $\gamma_h$ | 0.027 | 22.74 | <1.0x10<sup>-10</sup> |
| | **Wake v Anes**<br>**Cross-area**<br>**CL ⇔ LIP<sub>s</sub>**<br>N<sub>Wake</sub> = 5282<br>N<sub>Anes</sub> = 10478<br>DF = 15466 | $\delta$ | 0.027 | 24.40 | <1.0x10<sup>-10</sup> |
| | | $\theta$ | 0.123 | 83.19 | <1.0x10<sup>-10</sup> |
| | | $\alpha$ | 0.150 | 89.64 | <1.0x10<sup>-10</sup> |
| | | $\beta$ | 0.031 | 26.12 | <1.0x10<sup>-10</sup> |
| | | $\gamma_l$ | 0.058 | 54.91 | <1.0x10<sup>-10</sup> |
| | | $\gamma_h$ | 0.031 | 34.47 | <1.0x10<sup>-10</sup> |
| | **Wake v Anes**<br>**Cross-area**<br>**CL ⇔ FEF<sub>D</sub>**<br>N<sub>Wake</sub> = 3561<br>N<sub>Anes</sub> = 5675<br>DF = 9074 | $\delta$ | -0.001 | -0.612 | 0.541 |
| | | $\theta$ | 0.086 | 48.14 | <1.0x10<sup>-10</sup> |
| | | $\alpha$ | 0.086 | 47.65 | <1.0x10<sup>-10</sup> |
| | | $\beta$ | 0.040 | 49.62 | <1.0x10<sup>-10</sup> |
| | | $\gamma_l$ | 0.054 | 58.00 | <1.0x10<sup>-10</sup> |
| | | $\gamma_h$ | 0.042 | 45.84 | <1.0x10<sup>-10</sup> |
| | **Wake v Anes**<br>**Cross-area**<br>**CL ⇔ LIP<sub>D</sub>**<br>N<sub>Wake</sub> = 5806<br>N<sub>Anes</sub> = 9581<br>DF = 15029 | $\delta$ | 0.024 | 21.69 | <1.0x10<sup>-10</sup> |
| | | $\theta$ | 0.147 | 94.76 | <1.0x10<sup>-10</sup> |
| | | $\alpha$ | 0.161 | 102.90 | <1.0x10<sup>-10</sup> |
| | | $\beta$ | 0.042 | 39.74 | <1.0x10<sup>-10</sup> |
| | | $\gamma_l$ | 0.072 | 60.78 | <1.0x10<sup>-10</sup> |
| | | $\gamma_h$ | 0.022 | 19.19 | <1.0x10<sup>-10</sup> |

**Table S8.** Statistical results for cross-area thalamocortical spike-field coherence. Analyses with model 18 performed for: delta (δ) = 0-4 Hz; theta (θ) = 4-8 Hz; alpha (α) = 8-15 Hz; beta (β) = 15-30 Hz; low gamma (γ_l_) = 30-60 Hz; and high gamma (γ_h_) = 60-90 Hz. Spike-field coherence between CL spikes and cortical LFPs in superficial (S) or deep (D) layers of FEF or LIP. Reported statistics are the slope (β_1_), F statistic (F) and Holm’s adjusted p-value (p_adj_) for the parameter of interest (β_1_*State). Significant effects (p < 0.05) show frequency bands where coherence is significantly different for wakefulness relative to anesthesia.

| M18 | Data | Band | $\beta_1$ | F | $p_{adj}$ |
| --- | --- | --- | --- | --- | --- |
| spikeFC $\sim \beta_0 + \beta_1 * \text{State} + + \gamma_{\text{neuron}}^*(1)$ | **Wake v Anes**<br>**Cross-area**<br>**CL ↔ FEF<sub>s</sub>**<br>N<sub>Wake</sub> = 95<br>N<sub>Anes</sub> = 154<br>DF = 496.63 | $\delta$ | 0.021 | 47.95 | <1.0x10<sup>-10</sup> |
| | | $\theta$ | 0.029 | 154.65 | <1.0x10<sup>-10</sup> |
| | | $\alpha$ | 0.030 | 167.69 | <1.0x10<sup>-10</sup> |
| | | $\beta$ | 0.024 | 156.47 | <1.0x10<sup>-10</sup> |
| | | $\gamma_l$ | 0.018 | 112.75 | <1.0x10<sup>-10</sup> |
| | | $\gamma_h$ | 0.015 | 85.46 | <1.0x10<sup>-10</sup> |
| | **Wake v Anes**<br>**Cross-area**<br>**CL ↔ LIP<sub>s</sub>**<br>N<sub>Wake</sub> = 93<br>N<sub>Anes</sub> = 155<br>DF = 682.34 | $\delta$ | 0.029 | 135.27 | <1.0x10<sup>-10</sup> |
| | | $\theta$ | 0.028 | 159.64 | <1.0x10<sup>-10</sup> |
| | | $\alpha$ | 0.028 | 223.56 | <1.0x10<sup>-10</sup> |
| | | $\beta$ | 0.023 | 228.67 | <1.0x10<sup>-10</sup> |
| | | $\gamma_l$ | 0.018 | 188.59 | <1.0x10<sup>-10</sup> |
| | | $\gamma_h$ | 0.018 | 199.69 | <1.0x10<sup>-10</sup> |
| | **Wake v Anes**<br>**Cross-area**<br>**CL ↔ FEF<sub>D</sub>**<br>N<sub>Wake</sub> = 94<br>N<sub>Anes</sub> = 130<br>DF = 616.41 | $\delta$ | 0.024 | 67.50 | <1.0x10<sup>-10</sup> |
| | | $\theta$ | 0.028 | 171.18 | <1.0x10<sup>-10</sup> |
| | | $\alpha$ | 0.025 | 144.29 | <1.0x10<sup>-10</sup> |
| | | $\beta$ | 0.024 | 181.38 | <1.0x10<sup>-10</sup> |
| | | $\gamma_l$ | 0.021 | 181.10 | <1.0x10<sup>-10</sup> |
| | | $\gamma_h$ | 0.021 | 247.87 | <1.0x10<sup>-10</sup> |
| | **Wake v Anes**<br>**Cross-area**<br>**CL ↔ LIP<sub>D</sub>**<br>N<sub>Wake</sub> = 93<br>N<sub>Anes</sub> = 155<br>DF = 384.64 | $\delta$ | 0.035 | 125.98 | <1.0x10<sup>-10</sup> |
| | | $\theta$ | 0.032 | 181.38 | <1.0x10<sup>-10</sup> |
| | | $\alpha$ | 0.032 | 183.38 | <1.0x10<sup>-10</sup> |
| | | $\beta$ | 0.029 | 220.95 | <1.0x10<sup>-10</sup> |
| | | $\gamma_l$ | 0.024 | 189.16 | <1.0x10<sup>-10</sup> |
| | | $\gamma_h$ | 0.022 | 171.24 | <1.0x10<sup>-10</sup> |

**Table S9.**
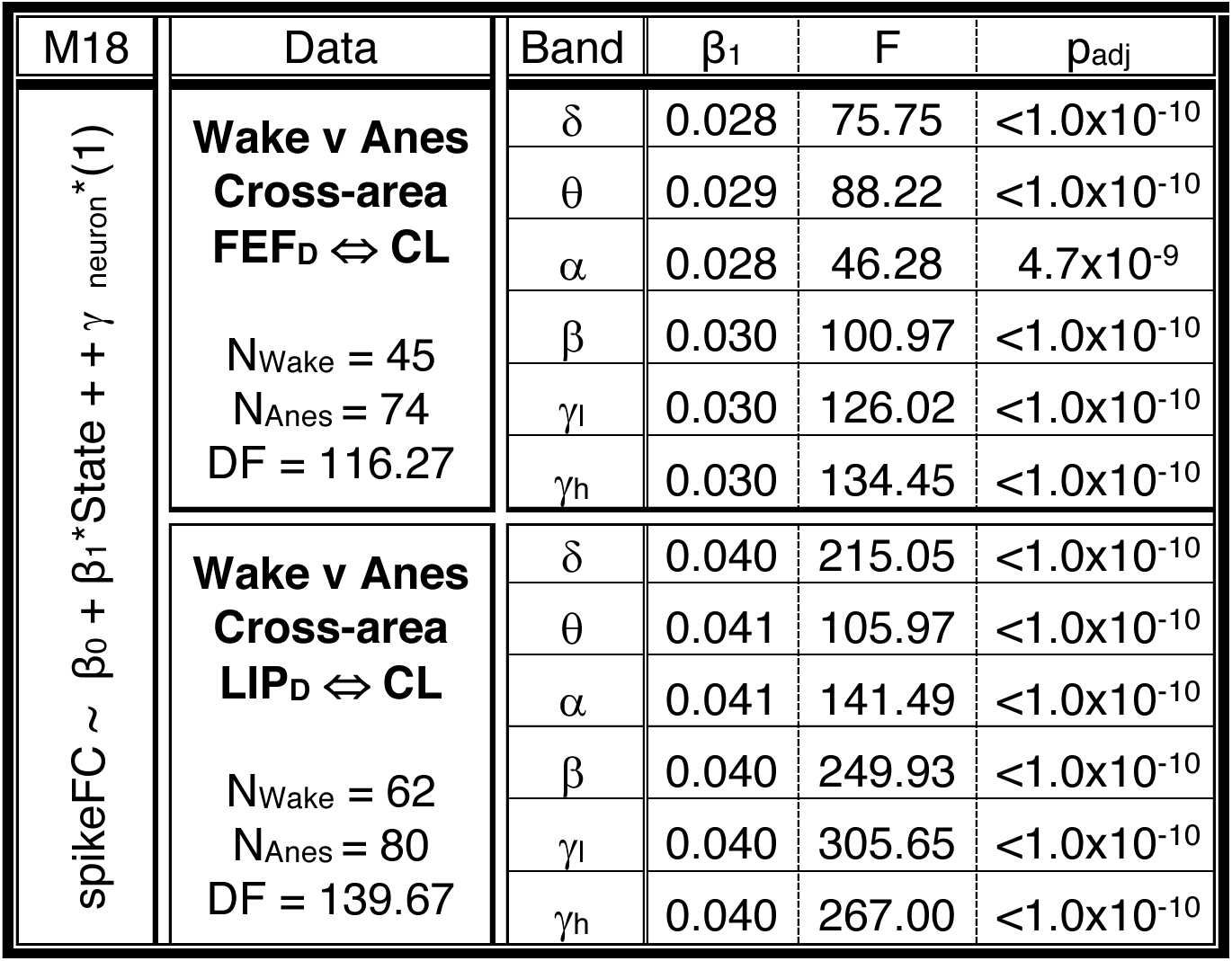
Statistical results for cross-area corticothalamic spike-field coherence. Analyses with model 18 performed for: delta (δ) = 0-4 Hz; theta (θ) = 4-8 Hz; alpha (α) = 8-15 Hz; beta (β) = 15-30 Hz; low gamma (γ_l_) = 30-60 Hz; and high gamma (γ_h_) = 60-90 Hz. Spike-field coherence between cortical spikes and CL LFPs. Reported statistics are the slope (β_1_), F statistic (F) and Holm’s adjusted p-value (p_adj_) for the parameter of interest (β_1_*State). Significant effects (p < 0.05) show frequency bands where coherence is significantly different for wakefulness relative to anesthesia.

| M18 | Data | Band | $\beta_1$ | F | $p_{adj}$ |
| --- | --- | --- | --- | --- | --- |
| $\text{spikeFC} \sim \beta_0 + \beta_1 * \text{State} + + \gamma_{\text{neuron}} * (1)$ | **Wake v Anes**<br>**Cross-area**<br>**FEF<sub>D</sub> ↔ CL**<br>$N_{\text{Wake}} = 45$<br>$N_{\text{Anes}} = 74$<br>$DF = 116.27$ | $\delta$ | 0.028 | 75.75 | $<1.0 \times 10^{-10}$ |
| | | $\theta$ | 0.029 | 88.22 | $<1.0 \times 10^{-10}$ |
| | | $\alpha$ | 0.028 | 46.28 | $4.7 \times 10^{-9}$ |
| | | $\beta$ | 0.030 | 100.97 | $<1.0 \times 10^{-10}$ |
| | | $\gamma_l$ | 0.030 | 126.02 | $<1.0 \times 10^{-10}$ |
| | | $\gamma_h$ | 0.030 | 134.45 | $<1.0 \times 10^{-10}$ |
| | **Wake v Anes**<br>**Cross-area**<br>**LIP<sub>D</sub> ↔ CL**<br>$N_{\text{Wake}} = 62$<br>$N_{\text{Anes}} = 80$<br>$DF = 139.67$ | $\delta$ | 0.040 | 215.05 | $<1.0 \times 10^{-10}$ |
| | | $\theta$ | 0.041 | 105.97 | $<1.0 \times 10^{-10}$ |
| | | $\alpha$ | 0.041 | 141.49 | $<1.0 \times 10^{-10}$ |
| | | $\beta$ | 0.040 | 249.93 | $<1.0 \times 10^{-10}$ |
| | | $\gamma_l$ | 0.040 | 305.65 | $<1.0 \times 10^{-10}$ |
| | | $\gamma_h$ | 0.040 | 267.00 | $<1.0 \times 10^{-10}$ |

**Table S10.** Statistical results for cross-area corticocortical coherence during effective and ineffective stimulations. Analyses with model 21 performed for: delta (δ) = 0-4 Hz; theta (θ) = 4-8 Hz; alpha (α) = 8-15 Hz; beta (β) = 15-30 Hz; low gamma (γ_l_) = 30-60 Hz; and high gamma (γ_h_) = 60-90 Hz for different pairs of contacts between FEF and LIP (Deep (D), Superficial (S), and Middle (M) layers). Reported statistics are the slope (β_1_), T statistic (T) and Holm’s adjusted p-value (p_adj_) for the parameter of interest (β_1_*StimEffect). Significant interactions (p < 0.05) show frequency bands where changes induced by stimulation are significantly different for effective relative to ineffective stimulations.

| M21 | Data | Band | $\beta_1$ | T | $p_{adj}$ |
| --- | --- | --- | --- | --- | --- |
| CDiff (stim – pre) $\sim \beta_0 + \beta_1 * \text{StimEffect} + \beta_2 * \text{DoseCode} + \beta_3 * \text{Anes}$ | **Stimulation Cross-area corticocortical LIP<sub>s</sub> ↔ FEF<sub>s/M</sub>**<br>N<sub>Effect</sub> = 1258<br>N<sub>Ineffect</sub> = 3397<br>DF = 2794 | $\delta$ | -0.021 | -4.05 | $8.6 \times 10^{-4}$ |
| | | $\theta$ | 0.002 | 0.49 | 1.000 |
| | | $\alpha$ | 0.026 | 6.87 | $1.5 \times 10^{-10}$ |
| | | $\beta$ | 0.002 | 0.83 | 1.000 |
| | | $\gamma_l$ | 0.012 | 4.45 | $1.5 \times 10^{-4}$ |
| | | $\gamma_h$ | 0.008 | 3.03 | 0.027 |
| | **Stimulation Cross-area corticocortical FEF<sub>D</sub> ↔ LIP<sub>s</sub>**<br>N<sub>Effect</sub> = 598<br>N<sub>Ineffect</sub> = 1974<br>DF = 1613 | $\delta$ | -0.026 | -2.92 | 0.032 |
| | | $\theta$ | 0.009 | 1.18 | 1.000 |
| | | $\alpha$ | 0.027 | 3.97 | 0.001 |
| | | $\beta$ | -0.012 | -3.10 | 0.024 |
| | | $\gamma_l$ | -0.001 | -0.41 | 1.000 |
| | | $\gamma_h$ | -0.014 | -3.44 | 0.008 |
| | **Stimulation Cross-area corticocortical FEF<sub>D</sub> ↔ LIP<sub>D</sub>**<br>N<sub>Effect</sub> = 523<br>N<sub>Ineffect</sub> = 2094<br>DF = 1666 | $\delta$ | -0.019 | -2.14 | 0.226 |
| | | $\theta$ | -0.003 | -0.37 | 1.000 |
| | | $\alpha$ | 0.001 | 0.14 | 1.000 |
| | | $\beta$ | -0.011 | -3.01 | 0.027 |
| | | $\gamma_l$ | -0.009 | -2.47 | 0.109 |
| | | $\gamma_h$ | -0.013 | -3.36 | 0.010 |

